# Functional Exploration of African Colorectal Cancer Patients Using Personalised *Drosophila* Avatars

**DOI:** 10.64898/2026.03.26.714433

**Authors:** Festus Adeyimika Oyeniyi, Favour Ayomide Oladokun, Abigael Abosede Ajayi, Abdulwasiu Ibrahim, Ruth Seyi Aladeloye, Opeyemi Abigail Akinfe, Funmilayo Rosemary Oludaiye, Thomas G. Moens, Hammed A. Badmos, Amos Olalekan Abolaji, Ross Cagan

## Abstract

Colorectal cancer across sub-Saharan Africa presents a growing global health burden, with increasing cases and mortality linked to late diagnosis, limited healthcare access and lack of effective treatments. African patients typically present with aggressive disease marked by distinct genomic signatures, indicating the need for targeted treatment approaches. We integrated genetic modelling, phenotypic scoring, imaging and biochemical analysis to explore how mutations found in individual Nigerian colorectal cancer patients influence drug responsiveness. We used the data from Cancer Genome Atlas to identify mutation profiles specific to Nigerian patients. We then generated ten stable *Drosophila melanogaster* personalised patient avatar lines designed to model patient genomic profiles. This study focused on three lines; each line included oncogenic RAS plus targeting patient-specific variants. These models exhibited various phenotypes including altered larval size, gut size and reduced survival. Two of the three avatar lines showed improved survival, reduced hindgut proliferation zone expansion and restored redox balance after treatment with regorafenib and trametinib. Mirroring clinical patient responses, we found that response to therapy is dependent on the specific genetic profile of the tumour.

**Graphical Abstract:** 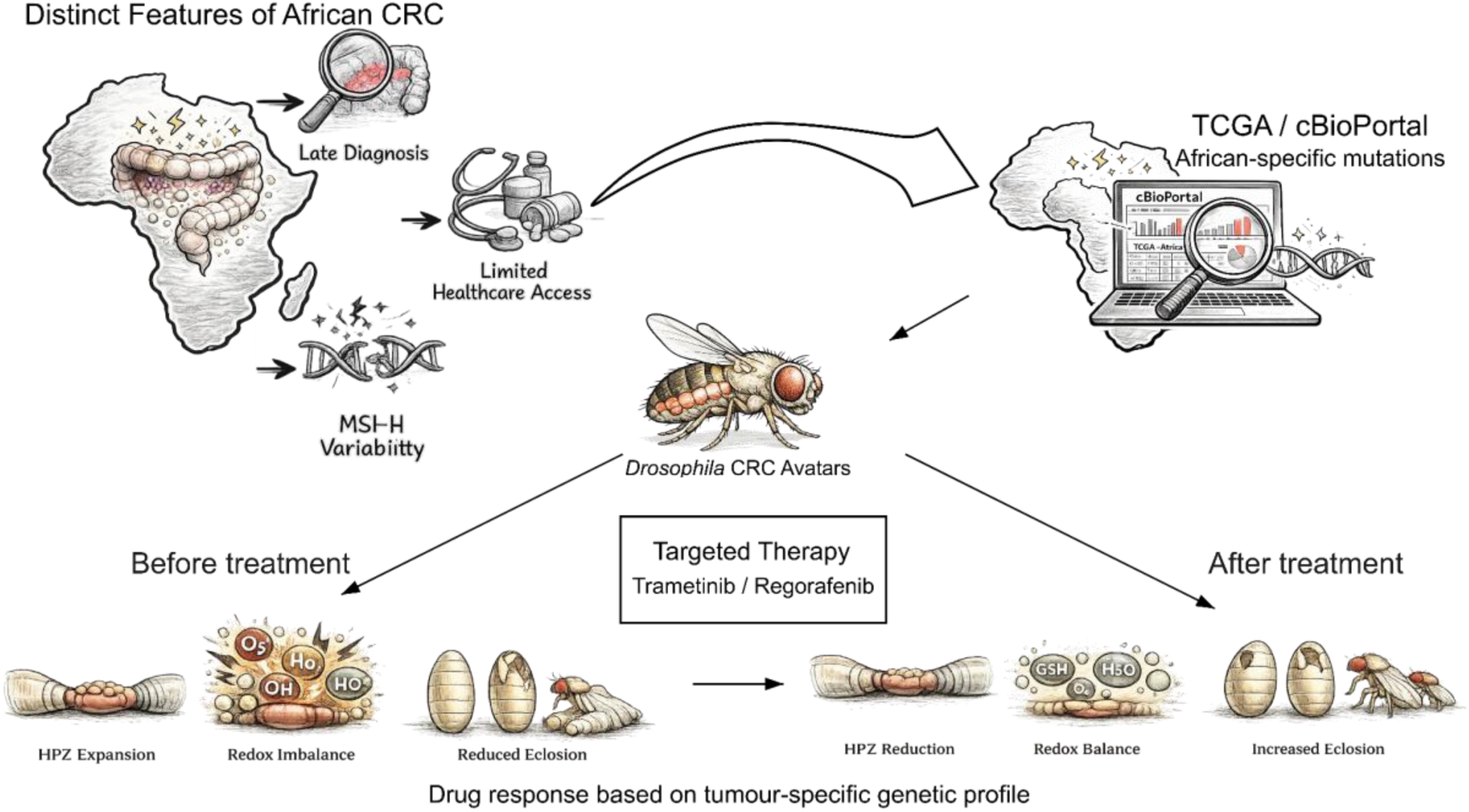

1. African colorectal cancer showed distinct mutation patterns that contribute to tumour heterogeneity.
2. Patient-derived *Drosophila* avatars were engineered using tumour-specific genetic mutations with key features of human colorectal cancer.
3. Treatment with targeted therapies showed responses patterned by tumour genotype.
4. Response patterns indicated the need for personalised for colorectal cancer therapies among diverse populations.

## Introduction

Colorectal Cancer (CRC) is a leading cause of cancer-associated deaths globally and the third most lethal cancer affecting men and women (Bray *et al*., 2024; Knapp *et al*., 2019). Approximately, 2 million new cases of CRC and 904,000 deaths were recorded in 2022 (Bray *et al*., 2024). High-Income Countries (HICs) have recorded significant progress in early diagnosis, prognosis and overall survival compared to African cohorts. However, in many Low- and Middle-Income Countries (LMICs), notably in sub-Saharan Africa, there is an alarming rise in CRC cases and deaths (Arnold *et al*., 2017). In Nigeria, for instance, CRC was previously considered rare. However, CRC incidence in Nigeria has recently risen to become the fourth most prevalent cancer type (Alatise *et al*., 2022). The high mortality rate from CRC in the African population is shaped by several factors such as late diagnosis and limited availability of treatments adapted to local context.

Advances in genomic studies continue to reveal population-specific biological differences that underlie how CRC develops and progresses in different populations. In a study of over 46,000 colorectal adenocarcinoma cases, African patients presented with aggressive tumours marked by higher rates of oncogenic alterations in KRAS and PIK3CA, lower rates in APC/WNT especially in younger patients and lower Microsatellite Instability-High (MSI-H) rates compared to their European counterparts (Alatise *et al*., 2021; Myer *et al*., 2022). This evidence implies that effective CRC treatment strategies must account for ancestral and ethnic variations.

Current CRC treatments include surgery, immunotherapy, radiotherapy and chemotherapy, with standard treatment often centred on 5-Fluorouracil in combination with FOLFOX or FOLFIRI regimens (Fadlallah *et al*., 2024; Singh *et al*., 2024). The clinical integration of targeted agents like EGFR and VEGF inhibitors has further broadened CRC treatment options (Heinemann *et al*., 2014; F. Li *et al*., 2023). Despite considerable improvement in treatment outcomes, efficacy varies due to the underlying genetic profile of the tumour plus its local and whole-body context (Kiran *et al*., 2024). For example, EGFR inhibitors are ineffective in CRC patients with *KRAS* mutations, while conventional chemotherapy has limited effect on patients with *BRAF-*mutant tumours (Gong *et al*., 2016). Accounting for this CRC heterogeneity will likely require identifying biomarkers that account for differences in ancestry that, in turn, impact the genetic landscape of African CRC patients across the region.

One tool to address this whole-body complexity is *Drosophila*. Flies provide a strongly accessible genetic system, conserved signalling pathways and a short life cycle, making them ideal for high-throughput *in vivo* modelling of CRC through the introduction of patient-specific mutations (Bangi *et al*., 2016; Cagan *et al*., 2019). Recent work highlights its utility, particularly the availability and the development of patient-specific fly avatars to model human cancer genotypes and facilitate high-throughput drug screening (Badmos & Cagan, 2025; Bangi *et al*., 2019).

We previously published patient reports from a ‘fly-to-bedside’ clinical trial that used ‘personalised fly avatars’ to screen for bespoke drug cocktails, including that of CRC patients (*NCT02363647*) (Bangi *et al*., 2019). In this study, we used tumour genetic data from ten previously characterised West African patients with microsatellite-stable (MSS) CRC (Alatise *et al*., 2021) to generate ten ‘personalised fly avatar’ transgenic lines, one for each modelled patient. These models allowed hindgut-specific targeting of key cancer driver genes.

To assess treatment response, we treated the fly avatars with regorafenib, a clinically relevant multi-kinase inhibitor approved for use in CRC, or with the potent MEK inhibitor trametinib. Microsatellite Stable (MSS) CRC is generally less responsive to immunotherapy, making pathway-targeted agents an especially important option (Gandini *et al*., 2023). In particular, the MAPK/ERK pathway targeted by trametinib has a central role in CRC progression and treatment resistance, and the kinase pathways targeted by regorafenib are also implicated in these processes (Q. Li *et al*., 2024).

A key property of CRC is redox imbalance, which is linked to cancer progression and resistance to therapy. Here, we use our Nigerian CRC avatar lines to assess levels of redox, including thiol levels and how these properties impact drug response. By combining genetic modelling with phenotypic scoring, imaging and biochemical assays, we show how *Drosophila* avatar lines carrying African patient-derived CRC mutant profiles respond to treatment and how these responses impact aspects of tumour progression.

## Results

### Model building and validation of *Drosophila* avatars engineered with Nigerian-derived CRC mutations

CRC in Nigerian patients exhibit significantly higher rates of *KRAS* alterations compared to Western cohorts (76.1% vs. 59.6%) (Alatise *et al*., 2021) - a tumour type for which there are few durable therapeutic options. We therefore focused on building avatars to model and explore this group of KRAS-driven CRC patients. Using the MSK-IMPACT sequencing panel analysis of 64 Nigerian CRC patients (Alatise *et al*., 2021) as reported in cBioPortal (cbioportal.org), we selected 10 KRAS mutant CRC patients for modelling. KRAS is commonly found with variants in APC and TP53 and the ten tumours we selected also contained mutations in APC (9/10) and TP53 (8/10; **Table 1**).

**Table 1.**
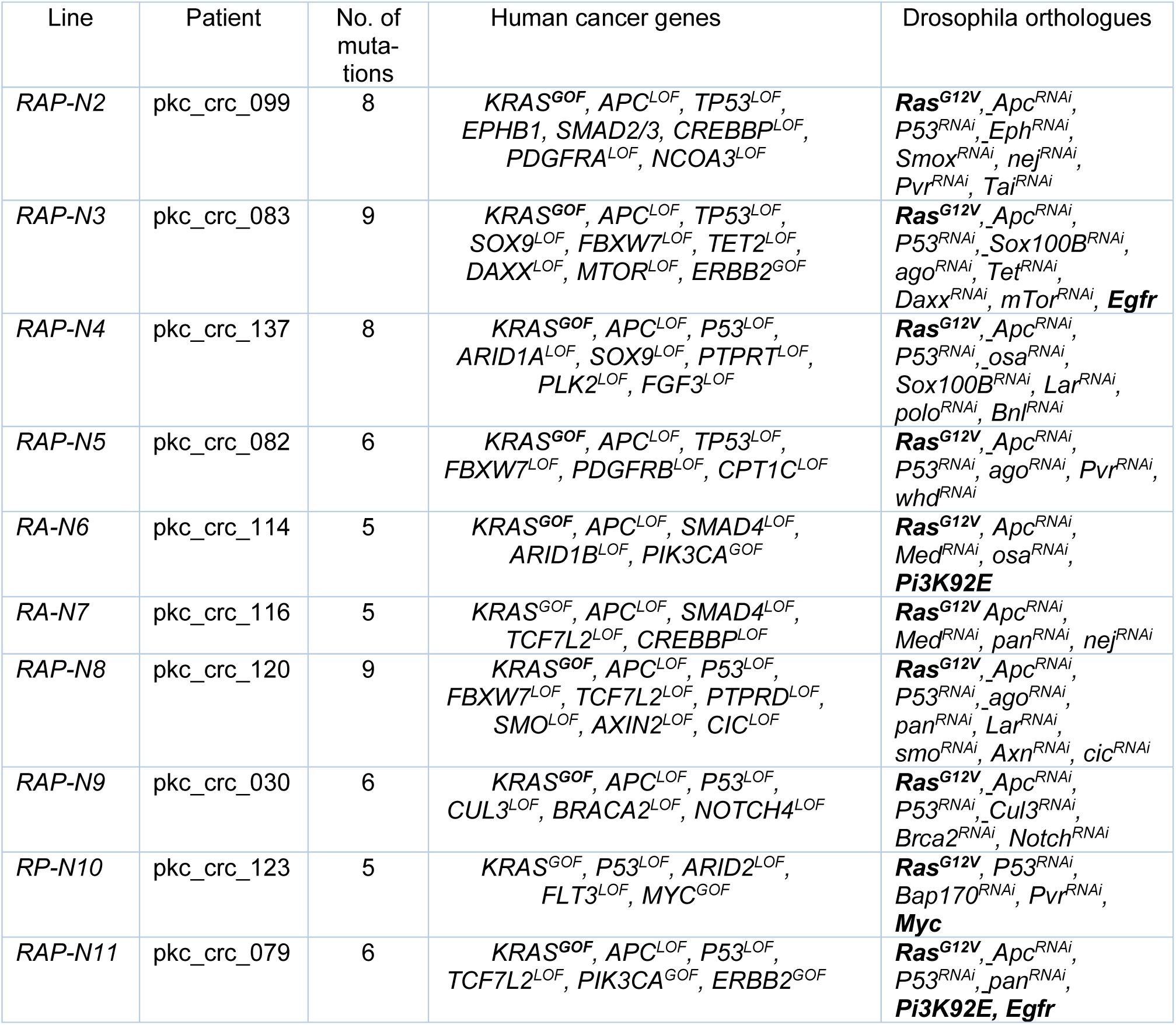

In addition to KRAS, APC and TP53 (*Drosophila Ras, Apc, p53*, referred to here as *RAP*), patients’ samples included a larger set of patient-specific mutations in known cancer-related genes. We targeted these additional mutations based on whether the mutation was a known oncogene or tumour suppressor, whether the mutation was predicted to be pathogenic or benign, penetrance of the mutations, allele frequency of the mutation in other CRC patients, and whether the gene is well-conserved in *Drosophila*. The oncogenes were modelled *via* overexpression, while tumour suppressors were modelled using shRNA-mediated knockdown. A summary of the selected mutations is shown in **Table 1** and construction details of the transgenic lines are found in **Supplementary Tables 1, 2 & 3**.

In brief, to model the effects of the selected mutations in *Drosophila*, we used the *GAL4/UAS* system, which allows for tissue-specific expression of *UAS*-directed transgenes in specific tissues. Using our previously reported transgene approach to multi-gene targeting (Bangi *et al*., 2019), we used overexpression (oncogenes) and RNA-interference-mediated knockdown (RNAi, tumour suppressors) to target a large cohort of cancer genes in a single transgene. For example, we combined activated *Drosophila RAS* (*Ras85D^G12V^*), RNA-interference-based knockdown of *APC* (*Apc^RNAi^*); and knockdown of *TP53* (*p53^RNAi^*) plus additional patient-specific gene targets to generate ten unique transgenic lines. We refer to these models as *RAP-N2* to *RAP-N11* (*Ras85D, Apc, p53* [*RAP*] or a subset of these [*RA*, *RP*]); reference numbers of the specific patients modelled are also found in **Table 1**. Our purpose was to capture and survey functional details of each patient beyond core mutations such as *Ras*.

Overexpression/knockdown of large sets of cancer driver genes in the *Drosophila* hindgut leads to tumour induction and developmental lethality (Bangi *et al*., 2016; Datta *et al*., 2023). We therefore used standard genetic crosses to combine our patient-specific lines with the hindgut specific *byn-GAL4* driver (abbreviated *byn>RAP-N\**; **Figure 1A**) as well as a temperature-sensitive *GAL80* allele (McGuire *et al*., 2003) that represses *GAL4* at low temperatures; the result is both spatial and temporal control over transgene expression. We also included a *UAS-GFP* reporter to aid visualisation of the transformed hindgut tissue.

**Figure 1:**
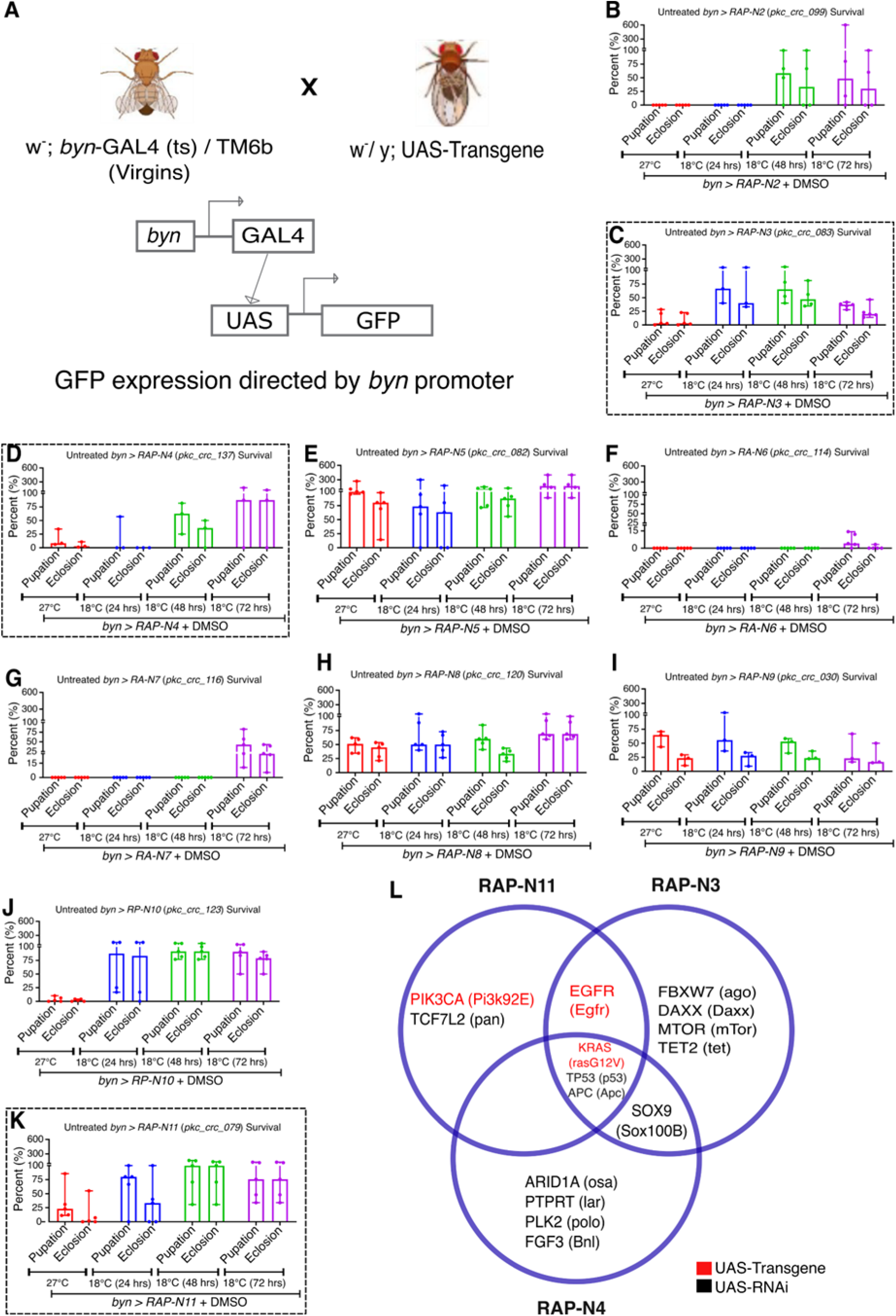
Model building, validation and phenotypic characterisation show. **(A)** Crosses of Nigerian Avatar lines with w^−^;; *byn-GAL4*, pTub-GAL80TS, UAS-GFP/TM6b GAL80 to express the tumour in the hindgut with a GFP reporter **(B-K).** Temperature calibration of ten CRC patient-derived avatars (*byn>RAP-N2* to *byn>RAP-N11)* (n=3 to 5 samples per group). The survival of *byn>RAP-N3, byn>RAP-N4 and byn>RAP-N11* are highlighted by the broken line squares **(C, D & K).** (**L**) a Venn diagram of overlapping mutations with their *Drosophila* orthologs in the highlighted three models. In (B-K), data are presented as median ± range. *p* < 0.05 (two-way ANOVA with Tukey’s post-hoc test).

We examined the baseline impact of tumour progression on the avatars at different temperatures (27 °C throughout development or the non-GAL4-permissive 18 °C for 24, 48 and 72 hours before shifting to 27 °C) to identify the optimal condition for transgene expression and tumour formation as assessed by lethality. Six of ten avatars (*byn>RAP-N3, N4, N8, N9, N11* and *byn>RP-N10*) exhibited reduced pupation and eclosion (survival to adulthood) at 27 °C throughout when compared to embryos kept initially at 18°C for 24, 48 and 72 hours, respectively, then switched to 27 °C. While two of ten *(byn>RA-N6 and N7*) showed increased lethality at 18°C for 72 hours. *byn>RAP-N2* showed increased lethality at 18°C for 48 hours, while *byn>RAP-N5* showed no lethality in all conditions (Figures **1B-K**).

We selected the most lethal condition specific to each fly line as a baseline for further studies, as it effectively modelled the lethality associated with the Nigerian-specific CRC mutational profiles. Of note, we focused on larvae due to their aggressive feeding behaviour and are more accessible to administering drugs (Bangi *et al*., 2016, 2019).

### Nigerian CRC patient-derived *Drosophila* avatars exhibited hindgut overgrowth

The mammalian and *Drosophila* intestines share several anatomical and physiological similarities (Jiang & Edgar, 2012). Targeting cancer genes to the hindgut can lead to gut hypertrophy, over-proliferation, loss of hindgut-specific markers and migration of cells to other sites in the fly (Bangi *et al*., 2016, 2019). To better understand how targeting these patient-specific genes can disrupt gut biology, we focused on three lines-*RAP-N3*, *RAP-N4*, and *RAP-N11*-with distinct genomic profiles, representing high (*RAP-N3*, *RAP-N4*) and lower (*RAP-N11*) genomic complexity (**Figure 1L**).

Compared to control flies expressing GFP alone (*byn>w^1118^)*, we observed significant expansion of the Hindgut Proliferation Zone (HPZ): *byn>RAP-N3*, *byn>RAP-N4* and *byn>RAP-N11* exhibited 123, 152 and 208% expansion in the HPZ, respectively, (**Figure 2B-E**). Further, we observed loss of *byn>GFP* signal in the HPZ of all three avatars, significantly in *byn*>*RAP-N4* guts (**Figure 2F**), consistent with a loss of *byn*-dependent cell identity.

**Figure 2:**
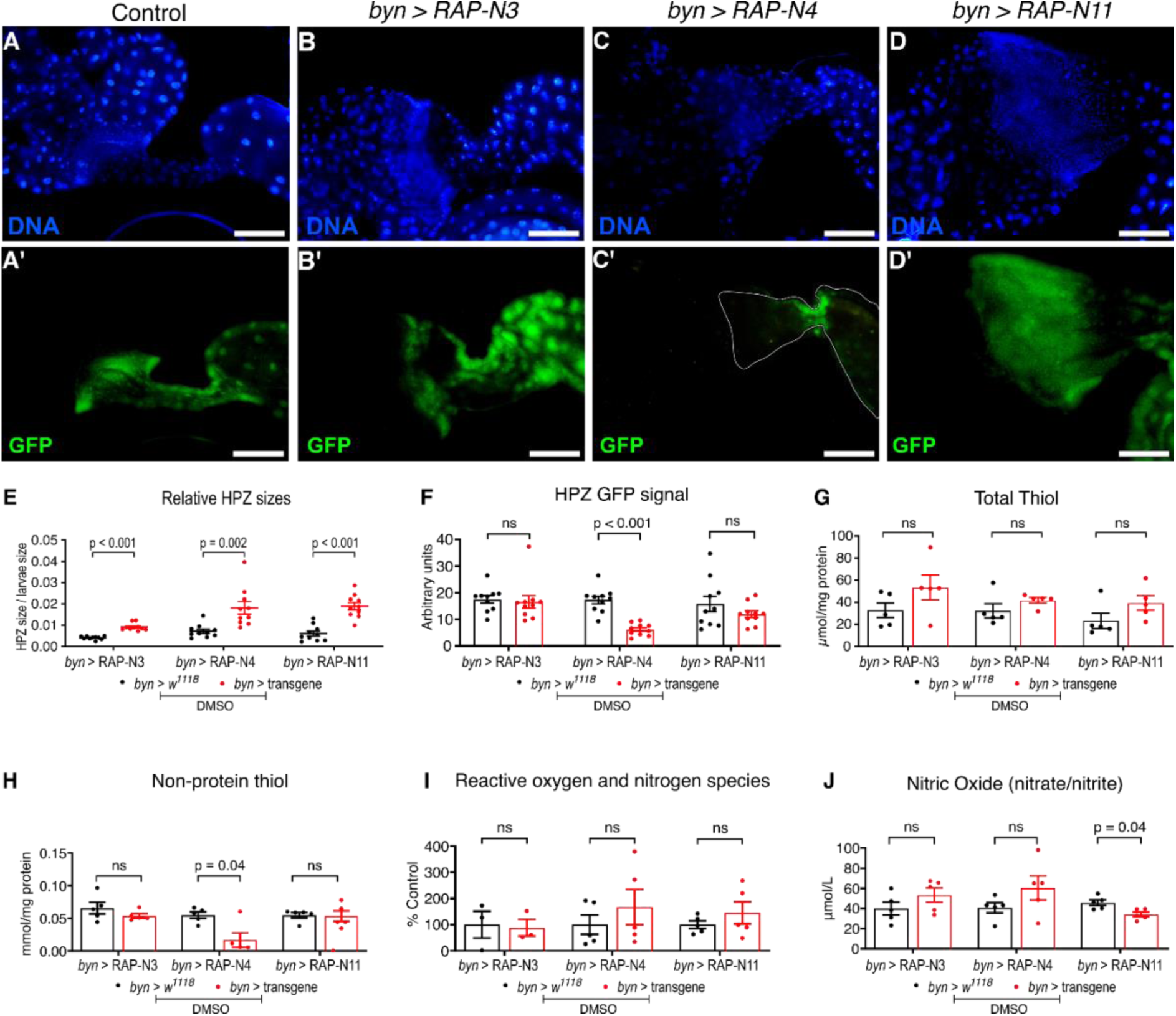
Characterisation of gut morphology of three patient-specific Nigerian-CRC fly Models: (A-D) GFP fluorescence imaging of larvae hindguts of *byn>RAP-N3, byn>RAP-N4 and byn>RAP-N11*. Scale bar: 100 μm. The images were processed using ImageJ (n=10 *Drosophila* hindgut for each group (E, F): Relative gut size (E) and (F) GFP fluorescence intensity of the hindgut proliferation zones (HPZ) of *byn>RAP-N3, byn>RAP-N4 and byn>RAP-N11* as compared to the control (n=10 *Drosophila* hindguts for each group) (G-H), antioxidant markers (I-J) and oxidative stress marker (n = 5 replicates per group). In (E-J), data are presented as mean ± SEM. p < 0.05 (unpaired two-tailed Student’s t-test).

We conclude that these three Nigerian avatar lines show important aspects of transformation including HPZ expansion and loss of at least one cell identity marker.

### Nigerian CRC patient-derived *Drosophila* avatars exhibited differential responses to targeted therapy

Given that the three Nigerian avatar lines showed clear features of transformation, we next tested their response to clinically relevant targeted CRC therapies. We mixed the drug into the larval food and assessed response to regorafenib (a multi-kinase inhibitor approved for CRC) and trametinib (a MEK1/2 inhibitor). At 5 days, untreated *byn>RAP-N4* and *byn>RAP-N11* showed marked reduction in larval sizes; however, regorafenib at 0.01 and 0.10 µM improved larval sizes for *byn>RAP-N4* compared to untreated control. Whereas regorafenib and trametinib-fed *byn>RAP-N3* and *byn>RAP-N11* showed no changes in larval sizes compared to untreated control (**Figures 3A-F**).

**Figure 3:**
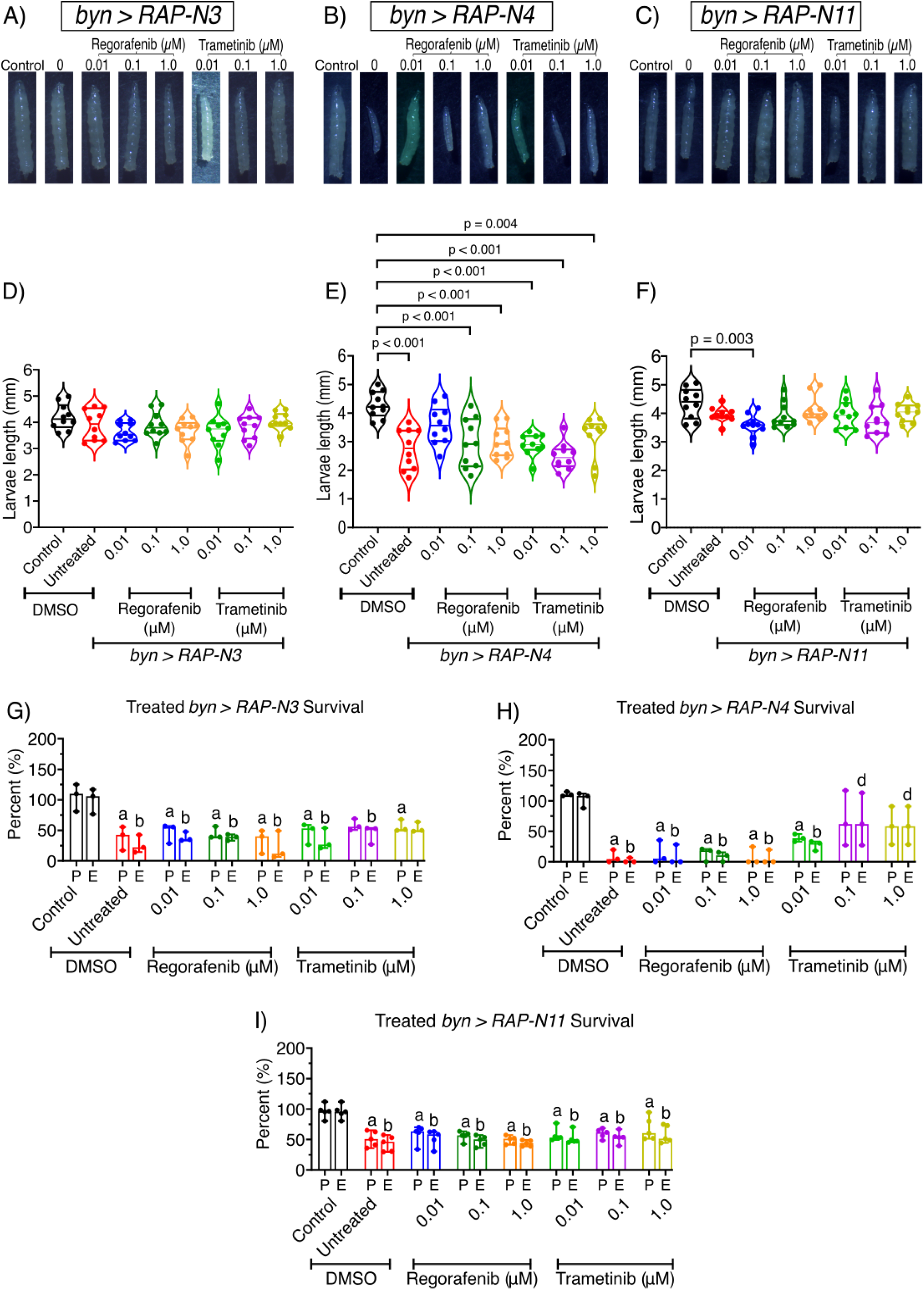
Effects of regorafenib and trametinib on larval sizes and survival rates across patient-specific Nigerian-CRC fly models. (A-F) larval size of *byn>RAP-N3*, *byn>RAP-N4* and *byn>RAP-N11* in response to regorafenib and trametinib at 0.01, 0.10 and 1.00 μM *(n = 10 larvae per group)*. (G-I) regorafenib and trametinib markedly improved pupation and eclosion rates of *byn>RAP-N3*, *byn>RAP-N4*, and *byn>RAP-N11* upon treatment with regorafenib and trametinib *(n = 3 to 5 sample replicates per group)*. (P = Pupation, E = Eclosion). (A–F) Mean ± SEM (ANOVA, Tukey’s); (G–I) median ± range. Significance (p < 0.05): **a**/**b**, reduction vs. control P/E; **d**, increase vs. untreated tumour E.

As shown in **Figure 1**, hindgut-specific expression of the transgenes reduced pupation and eclosion in all three lines, making survival to adulthood a useful readout of drug response. Using this assay, we tested regorafenib and trametinib as single agents at 0.01, 0.10 and 1.00 µM. Feeding *byn>RAP-N3* larvae regorafenib led to an increase in eclosion by 12% at 0.01 and 0.10 µM, while trametinib increased eclosion by 7, 18, and 27% at 0.01, 0.10, and 1.00 µM, respectively, as compared to untreated controls (**Figure 3G**). In *byn>RAP-N4* animals, trametinib caused significant rescue, increasing pupation by 31, 60, and 51% and eclosion by 25, 65, and 57% at 0.01, 0.10, and 1.00 µM, respectively, (**Figure 3H**). In contrast, *byn>RAP-N11* animals showed no improvement in survival with either drug at these concentrations (**Figure 3I**).

Together, these data indicate genotype-dependent sensitivity to regorafenib and trametinib in Nigerian CRC avatar lines. The resistance of *byn>RAP-N11* may reflect alterations in *Pi3K92E* (PIK3CA) and *pan* (TCF7L2), which could promote PI3K-AKT signalling and alter Wnt/β-catenin-dependent transcription, both of which have been linked to reduced drug sensitivity in CRC (Voutsadakis, 2025; Wang *et al*., 2024).

### Regorafenib and trametinib selectively improve hindgut architecture in Nigerian CRC patient-derived *Drosophila* avatars

To assess whether regorafenib and trametinib also improved tissue architecture, we examined their effects on the enlarged HPZ phenotype. Relative gut size was calculated by normalising HPZ area to larval area to account for differences in overall body size. Both drugs fed to *byn>RAP-N4* larvae selectively reduced HPZ expansion significantly at 0.01 μM regorafenib (*p* = 0.03) and trametinib (*p* = 0.04). Generally, 0.01 and 1.00 μM regorafenib-fed *byn>RAP-N4* larvae reduced HPZ by 49 and 29%, while 0.01 and 1.00 μM trametinib concentrations reduced HPZ by 38 and 47%, respectively, relative to untreated controls. In addition, *byn>RAP-N3* animals fed with 0.01 and 0.10 μM regorafenib reduced HPZ expansion by 15 and 16%, while trametinib reduced expansion by 26 and 23% at 0.10 and 1.00 μM, respectively. Further, regorafenib at 0.10 μM reduced *byn>RAP-N11* larvae HPZ by 10% while trametinib at 0.01, 0.10 and 1.00 μM caused a clear reduction in HPZ sizes by 27, 26 and 34%, respectively, in *byn>RAP-N11* when compared to the untreated control (**Figure 4A-X**).

**Figure 4:**
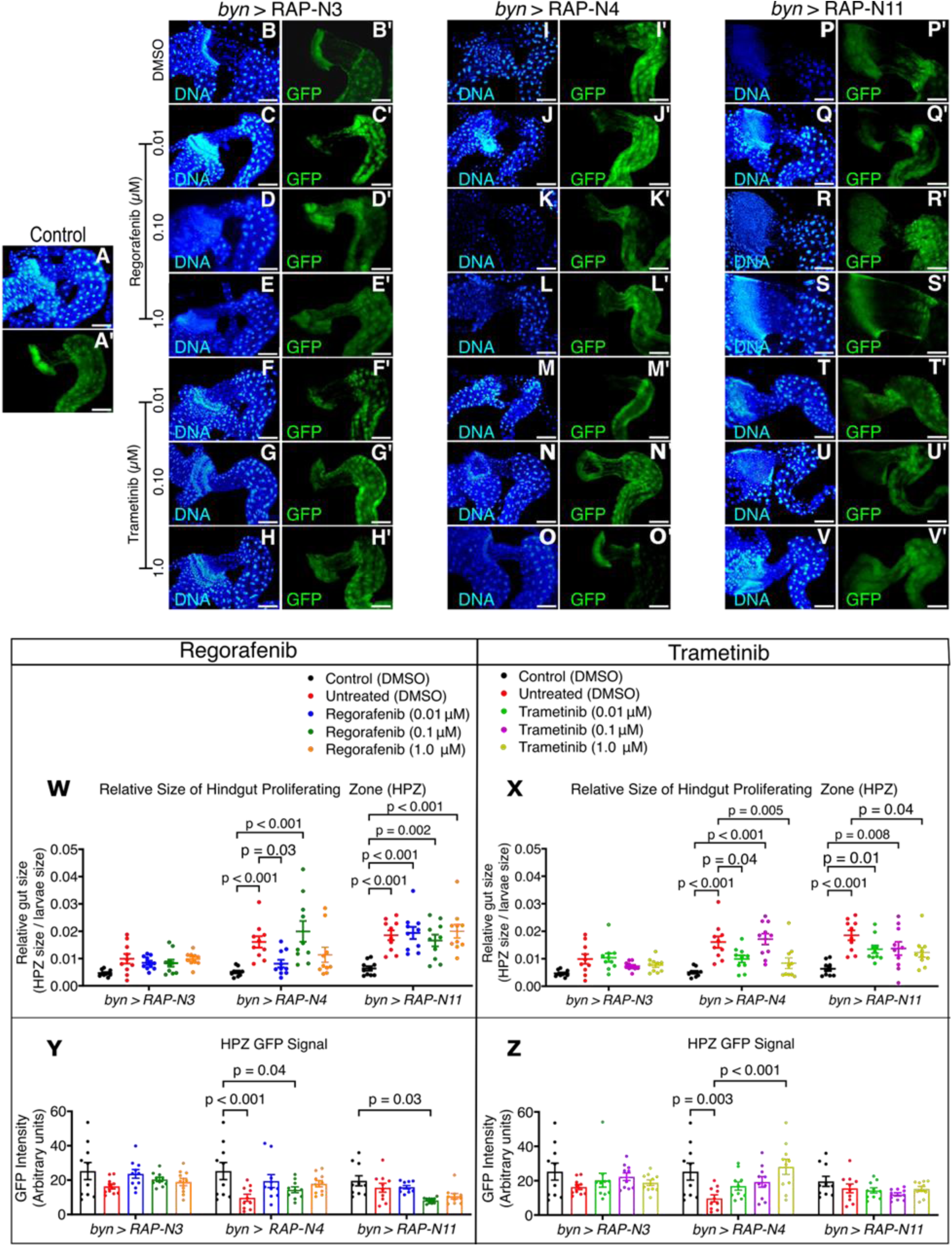
Therapeutic effects of regorafenib and trametinib on proliferating gut size and GFP Intensity; (**A-V**) Fluorescence micrographs of the hindgut proliferation zone (HPZ) showing the gut integrity of the models treated with regorafenib and trametinib. Scale bar: 100 μm (**W-X**) relative proliferating gut sizes of the models treated with regorafenib and trametinib *(n = 10 guts per group)*. **(Y-Z)** GFP intensity of the proliferating gut of the models treated with regorafenib and trametinib *(n = 10 guts per group).* In (**W-Z**), data are presented as mean ± SEM. p < 0.05 (one-way ANOVA with Tukey’s post-hoc test).

We next measured *byn>GFP* intensity as an additional readout of hindgut transformation. GFP intensity was restored in *byn>RAP-N3* and *byn>RAP-N4* after treatment with regorafenib or trametinib, but not in *byn>RAP-N11* (**Figure 4Y-Z**). Correlation analysis further showed that smaller HPZ size generally tracked with higher eclosion and higher GFP intensity, although the strength of these associations varied by genotype and treatment (**Supplementary Tables S4-S9**). Specifically, for the *byn>RAP-N3* avatar, correlation analysis revealed a strong inverse relationship between relative gut size and eclosion rates with both regorafenib (Table S4) and trametinib (Table S5) treatment, indicating that reduction in HPZ expansion is a strong predictor of improved adult survival in this genotype.

Together, these data indicate that regorafenib and trametinib partially reversed abnormal hindgut architecture in a genotype-dependent manner, notably in *byn>RAP-N4*. Only trametinib showed clear activity on *byn>RAP-N3* and *byn>RAP-N11* animals.

### Regorafenib and trametinib selectively inhibit *Drosophila* MAPK (dpERK) signalling in Nigerian CRC patient-derived *Drosophila* avatars

Phosphorylated Extracellular Signal-Regulated Kinase (Phospho-ERK), an activated form of ERK1/2, is a key kinase in the MAPK signalling involved in regulating cell proliferation, survival, differentiation and migration-processes that are frequently dysregulated in CRC (White *et al*., 2026).

To understand the expression of dpERK, we measured the mean signal of dpERK at the midgut-hindgut axis of the CRC avatars (**Figure 5).** Untreated *byn>RAP-N3* showed dpERK activation compared to control, which indicates active ERK signalling within the HPZ. However, animals fed with trametinib at all concentrations and those fed with regorafenib (0.10 and 1.00 μM) showed decreased dpErk intensity compared to the untreated animals, while 0.01 μM regorafenib showed a higher dpERK signal (**Figure 5A).** The untreated *byn>RAP-N4* group showed no clear difference in dpERK signal compared to control; however, trametinib at 0.10 and 1.00 μM caused a reduction in the dpErk intensity (**Figure 5B**). A significant increase was observed in dpERK signal of untreated *byn>RAP-N11* animals compared to control (*p* = 0.01). The treatment of *byn>RAP-N11* with regorafenib and trametinib led to a significant decrease across all concentrations compared to the untreated control (**Figure 5C**).

**Figure 5:**
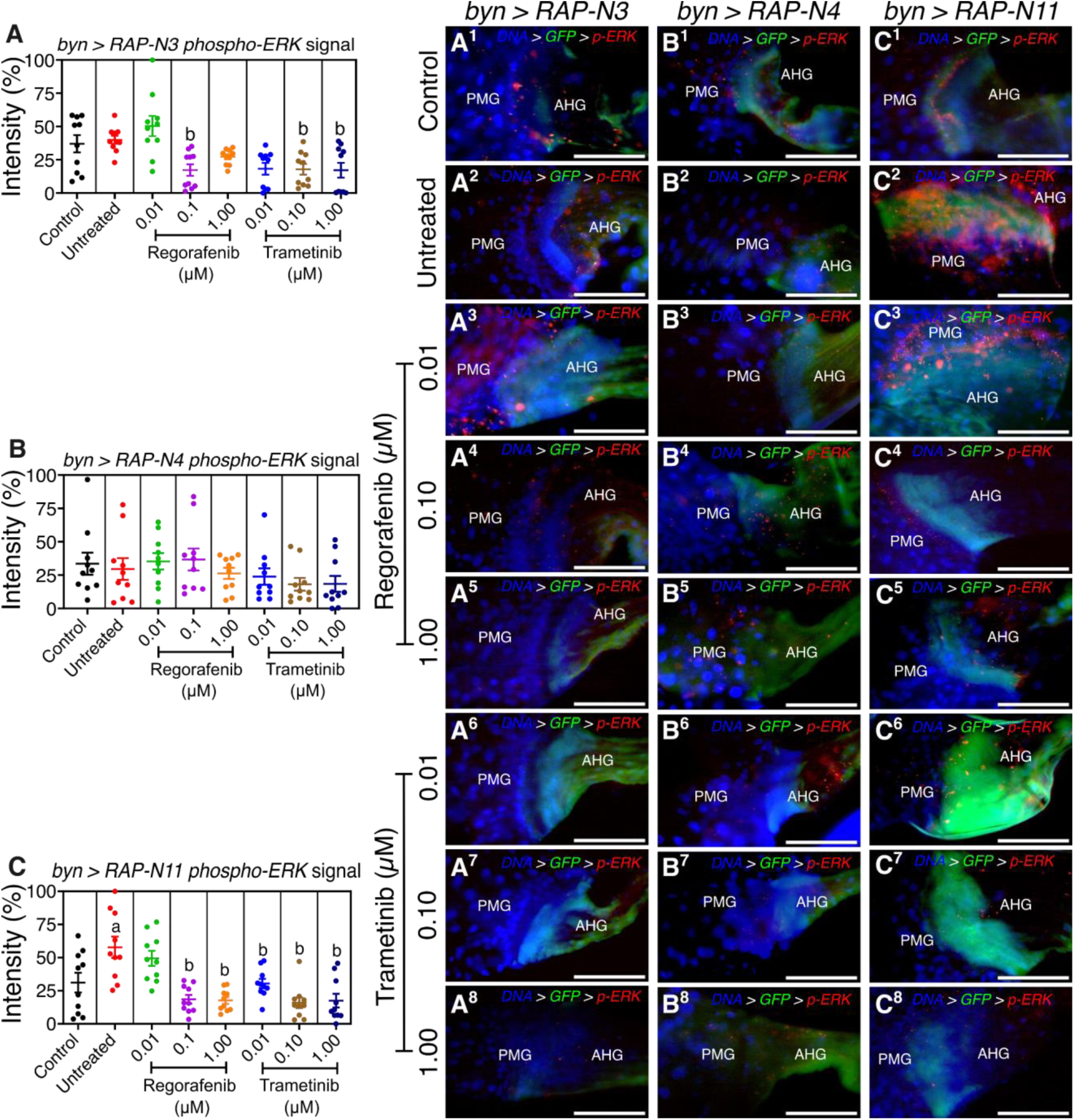
Effects of regorafenib and trametinib on the spatial distribution of *Drosophila* phospho-ERK (dpERK) intensity at the HPZ; (**A^1^-C^8^**) micrographs of the hindgut proliferation zone (HPZ) showing the dpERK signal of the models treated with regorafenib and trametinib. Scale bar: 100 μm (**A-C**) mean dpErk intensity along the midgut-hindgut axis of the avatar lines treated with regorafenib and trametinib *(n = 10 guts per group)*. In (**A-C**), data are presented as mean ± SEM. p < 0.05 (one-way ANOVA with Tukey’s post-hoc test). **AHG** = anterior hindgut; **PMG** = posterior midgut. Significance (p < 0.05): **a**, increase vs. control; **b**, decrease vs. untreated tumour.

Further, spatial analysis of dpERK intensity profile revealed a progressive increase in signal towards the distal hindgut across the CRC avatar lines (**Supplementary Figures S2–S4**). In Figure S2, dpERK intensity shows several localised peaks along the hindgut axis, with the most prominent signals appearing beyond ∼60 µm from the HPZ in the control and ∼80 µm in untreated *byn>RAP-N3*. A similar pattern was observed in regorafenib and trametinib fed *byn>RAP-N4* model (**Supplementary Figure S3**), where dpERK levels remain relatively low near the HPZ but increase further along the distal hindgut, with trametinib-treated samples showing pronounced peaks in the distal region while control and untreated groups maintained a low dpERK profile. In the *byn>RAP-N11* model (**Supplementary Figure S4**), dpERK intensity exhibits multiple distal peaks along the hindgut axis, particularly across untreated and 0.01 μM regorafenib treatments.

Together, these spatial profiles indicate that ERK signalling is not uniformly distributed along the hindgut but instead showed a consistent distal enrichment across the CRC avatar models.

### Untreated Nigerian CRC avatar lines exhibit mild redox imbalance

Oxidative stress is strongly associated with colorectal cancer, where increased Reactive Oxygen Species (ROS) can damage biomolecules and support tumour initiation and progression (Bardelčíková *et al*., 2023). As tumours develop, transformed cells often adapt by strengthening antioxidant systems, creating a redox state that supports survival and treatment resistance. To assess baseline redox status in the Nigerian CRC avatars, we measured total thiols, non-protein thiols, Reactive Oxygen and Nitrogen Species (RONS) and nitric oxide (measured as nitrite/nitrate level) in third instar larvae.

At baseline, total thiol level did not differ significantly from control in any of the three lines **(Figure 2G)**. Non-protein thiol levels, however, varied by genotype: *byn>RAP-N4* showed a significant reduction relative to control (p = 0.04), whereas *byn>RAP-N3* and *byn>RAP-N11* did not (Figure 2H). The nitric oxide and RONS readouts also differed across lines. The *byn>RAP-N4* and *byn>RAP-N11* showed marked increases in RONS, whereas an increase in nitric oxide was observed in *byn>RAP-N4* following a significant reduction in nitric oxide in *byn>RAP-N11* larvae compared to the control. In contrast, *byn>RAP-N3* showed no change in RONS level, while there was an increase in nitric oxide levels compared to control **(Figures 2I, J)**. Together, these data indicate that untreated Nigerian avatar lines do not share a single redox phenotype but show at most mild, genotype-dependent redox imbalance.

### Regorafenib and trametinib produced genotype-specific redox responses in Nigerian CRC avatar lines

To further affirm this observation, we evaluated the dynamics of antioxidant biomarkers across the fly models in response to regorafenib and trametinib. In the total thiol and non-protein thiol, we observed inconsistent patterns among the fly avatars; regorafenib fed *byn>RAP-N3* at 0.10 & 1.00 μM and *byn>RAP-N4* at 0.10 μM showed significant increases in total thiol level compared to untreated control. Also, trametinib-fed *byn>RAP-N3* and *byn>RAP-N4* animals showed no observable change compared to untreated controls. In addition, *byn>RAP-N11* showed a slight decrease in total thiol for both drugs compared to the untreated control (**Figures 6A-B**). For non-protein thiol levels, *byn>RAP-N3 and byn>RAP-N11* showed a slight reduction in non-protein thiol levels for both drugs relative to the untreated control, while *byn>RAP-N4* showed a significant increase at 1.00 μM of regorafenib and trametinib (**Figures 6C-D)**.

**Figure 6:**
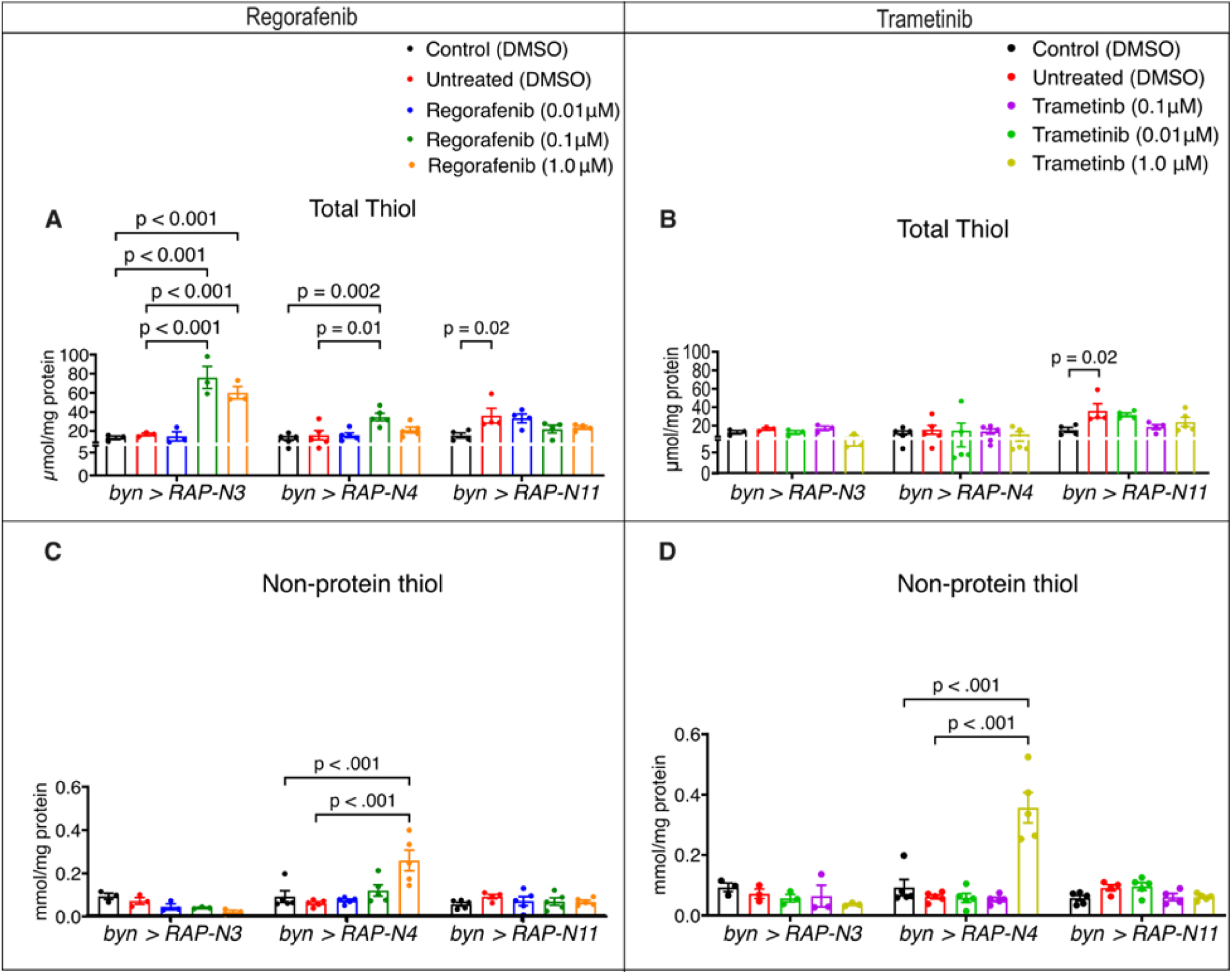
Therapeutic effects of regorafenib and trametinib on antioxidant markers. Total thiol and non-protein thiol in patient-specific *byn>RAP-N3*, *byn>RAP-N4* and *byn>RAP-N11*) avatar models. (A-B) Total thiol levels of larvae treated with regorafenib and trametinib *(n = 3 to 5 sample replicates per group).* (C-D) Non-protein thiol levels of larvae treated with regorafenib and trametinib *(n = 3 to 5 sample replicates per group)*. In (A-D), data are presented as mean ± SEM. p < 0.05 (one-way ANOVA with Tukey’s multiple-comparison test).

We evaluated the effects of regorafenib and trametinib on the redox environment of the three *byn>RAP-N* avatars, revealing that the *byn>RAP-N4* animals are uniquely predisposed to drug-induced oxidative stress (**Figure 7**). Remarkably, both drugs significantly induced RONS production specifically in *byn>RAP-N4* background (**Figures 7A, B**). For regorafenib-fed *byn>RAP-N4*, a concentration-dependent increase in RONS, with the highest concentrations (0.10 μM, *p* = 0.02 and 1.00 μM), was notable. Interestingly, trametinib induced a similar RONS spike at its lowest concentration (0.01 μM, *p* = 0.02) when compared to the control, following a non-linear trend as concentrations increased. Whereas *byn>RAP-N3* and *byn>RAP-N11* animals remained largely unresponsive or exhibited a decrease in RONS levels at higher concentrations. For example, 1.00 μM regorafenib led to a dramatic reduction in RONS level in *byn>RAP-N3,* while 0.01 μM trametinib reduced RONS levels in *byn>RAP-N11* compared to untreated animals.

**Figure 7:**
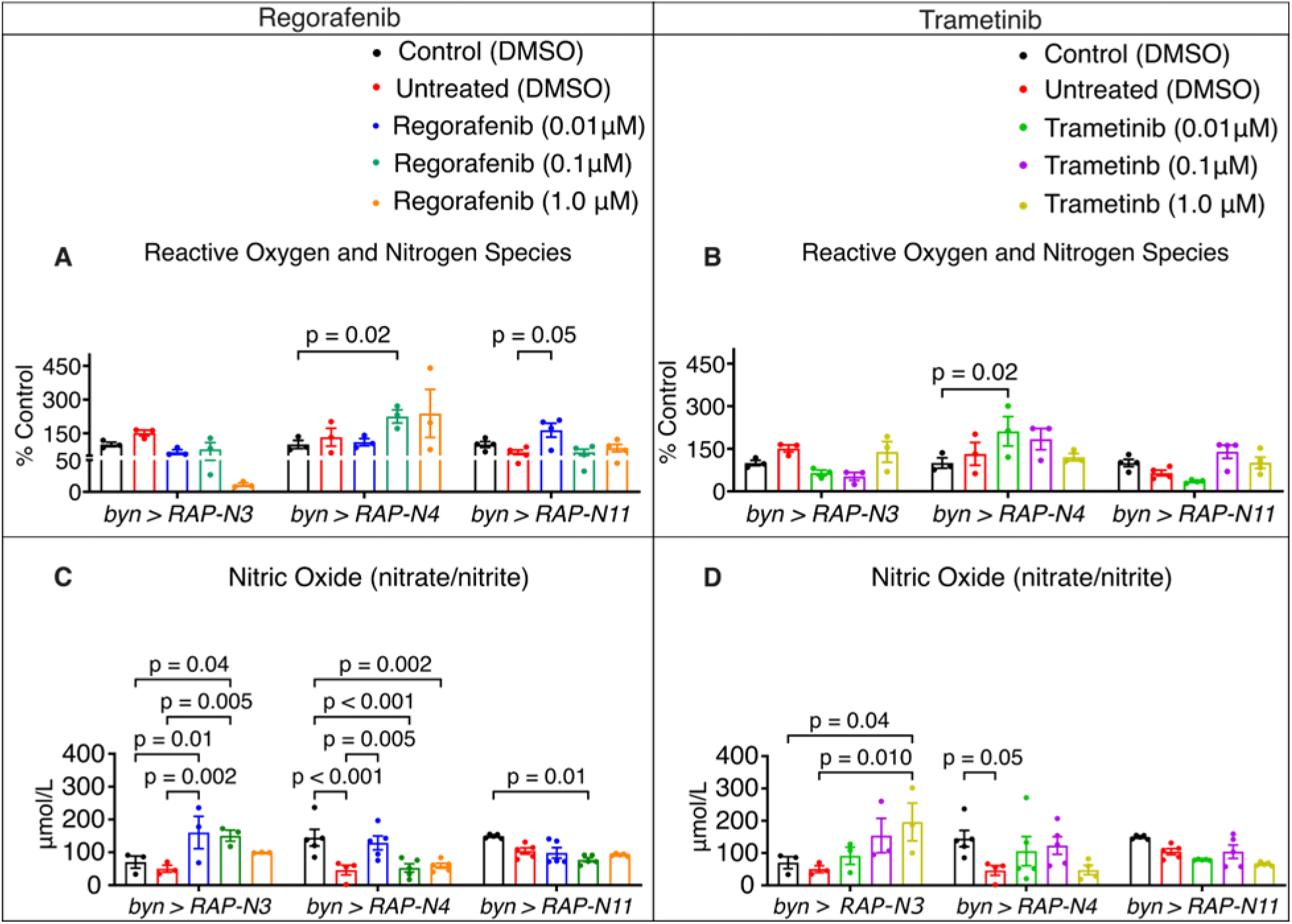
Therapeutic effect of regorafenib and trametinib on biomarkers of oxidative stress. RONS and NO in Patient-specific Nigerian-CRC Fly Models (*byn>RAP-N3*, *byn>RAP-N4* and *byn>RAP-N11*); (A-B) RONS levels of larvae treated with regorafenib and trametinib (*n = 3 to 5 sample replicates per group)*. (C-D) Nitric oxide of larvae treated with regorafenib and trametinib *(n = 3 to 5 sample replicates per group);* In (A-D), data are presented as mean ± SEM. p < 0.05 (one-way ANOVA with Tukey’s multiple-comparison test).

Parallel analysis of nitrosative stress showed a striking difference in how these avatars managed nitric oxide under drug intervention (**Figure 7C-D**). In *byn>RAP-N3* avatars, regorafenib and trametinib significantly induced a concentration-dependent increase in nitric oxide levels compared to the untreated animals (regorafenib: 0.01 μM, *p* = 0.02, 0.10 μM, *p* = 0.05 and trametinib: 1.00 μM, *p* = 0.01). Also, untreated *byn>RAP-N4* animals maintained significantly low nitric oxide level compared with the control (p < 0.001). Regorafenib at 0.01 μM rescued the suppressed nitric oxide levels, while higher concentrations show no observable increase.

In addition, trametinib showed marked improvement in nitric oxide level compared to untreated *byn>RAP-N4* animals. The *byn>RAP-N11* remained unresponsive to both drugs with respect to the restoration of nitric oxide levels. Furthermore, the *byn>RAP-N4* showed distinct correlation patterns, revealing a significant inverse correlation between relative gut size and nitric oxide (**Table S6**; r = -0.96, p = 0.04) for regorafenib treatment, as well as a strong correlation observed between the relative gut size and RONS in the trametinib treatment (**Table S7**; r = 0.91, *p* = 0.09). These genotype-dependent relationships reflect how population-specific genetic differences reshape redox status and therapeutic outcome.

### Mitochondrial metabolic rate and transcriptional response thioredoxin-2 (*Trx-2*) also varied significantly by genotype and drug type

Having established varying dynamics of biochemical markers of redox response in the models, we evaluated mitochondrial metabolic rate using the MTT assay and mRNA expression of thioredoxin-2 (*Trx-2*) as a molecular biomarker of CRC progression (Mollbrink *et al*., 2014). Mitochondrial metabolic rate and *Trx-2* mRNA expression showed that the three CRC-avatar genotypes responded differently to regorafenib and trametinib. The *byn>RAP-N3* showed an increased mitochondrial metabolic rate of both drugs, while *byn>RAP-N4* showed a significant increase at 0.1 μM trametinib compared to untreated control. However, *byn>RAP-N11* at 0.1 μM of both drugs indicated a decrease in mitochondrial metabolic rate as opposed to the untreated subset. This may suggest that the tumour genotypes activate mitochondrial pathways that boost metabolic output instead of undergoing drug-induced damage (**Figures 8A-B**).

**Figure 8:**
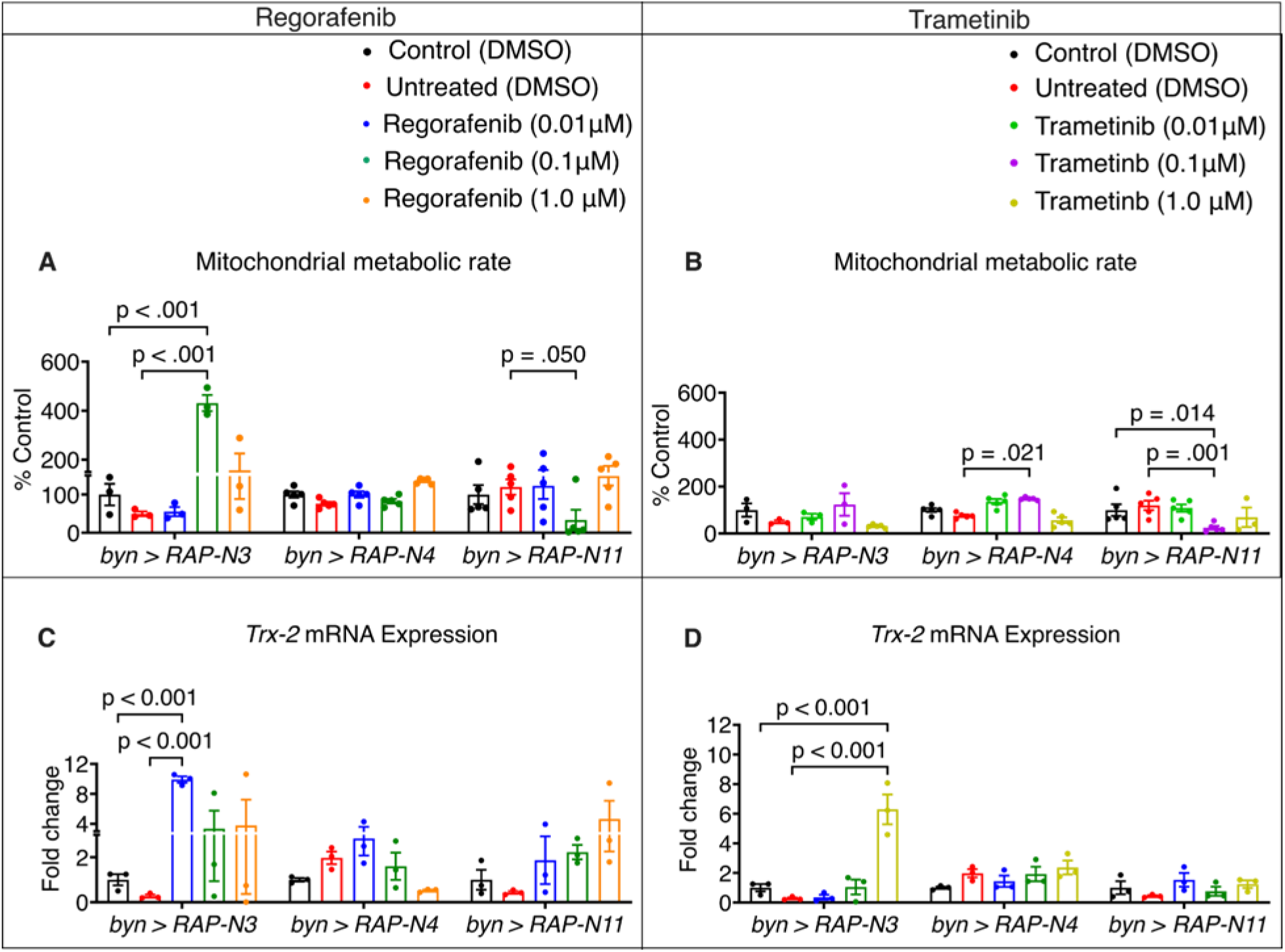
Effects of regorafenib and trametinib on mitochondrial metabolic activity and *Trx-2* mRNA expression. (A-B) Metabolic activity of larvae treated with regorafenib and trametinib *(n = 3 to 5 sample replicates per group)*. (C-D) *Trx-2* gene expression levels of larvae treated with regorafenib and trametinib *(n = 3 to 5 sample replicates per group)*. In (A-D), data are presented as mean ± SEM. p < 0.05 (one-way ANOVA with Tukey’s multiple-comparison test).

The increase in metabolic rate is accompanied by upregulation of mRNA expression of *Trx-2*, a mitochondrial protein that helps to maintain redox balance and protect metabolic enzymes while promoting drug resistance and cancer progression (Jovanović *et al*., 2022). Remarkably, regorafenib increased *Trx-2* mRNA levels in both *byn>RAP-N3* and *byn>RAP-N11* (**Figure 8C).** In contrast, trametinib reduced *Trx-2* mRNA expression in *byn>RAP-N4* (0.01 μM and 0.10 μM) but increased it in *byn>RAP-N3* and *byn>RAP-N11* (0.01 μM and 1.0 μM) when compared with their untreated controls (**Figure 8D)**.

## Discussion

Here, we report on patient-specific *Drosophila melanogaster* models that reflect the genetic variations in selected Nigerian CRC patients (Alatise *et al*., 2021). We used a previously developed technology used in a fly-to-bedside clinical trial (*NCT02363647*) (Bangi *et al*., 2019) to model individual Nigerian CRC patients. Our functional data support the use of patient-specific *Drosophila* avatars to define genotype-dependent drug responses in CRC (Bangi *et al*., 2019). Responses in fly tumour models often parallel those seen in patients, which supports their use as functional systems for testing targeted therapies in a personalised setting (Bangi *et al*., 2019, 2021).

In this study, regorafenib and trametinib improved larval size and reduced HPZ expansion in the *byn>RAP-N4* avatar, indicating that this genotype is treatment-responsive. This is notable because untreated *byn>RAP-N4* carries changes in growth-regulatory genes including *osa* (orthologue of human *ARID1A*) and *polo* (*PLK2*), which are linked to chromatin regulation, mitosis and tissue growth (Chapagai *et al*., 2025; Hao *et al*., 2025). The observed sensitivity of the lines to RAF and MEK inhibitions reflects fundamental biology conserved across species; of note, the drug concentrations used in our study should not be viewed as directly informing human dosing, given major interspecies differences in xenobiotic handling and metabolism (Rand *et al*., 2023). Nonetheless, the concentration-dependent effects on HPZ size and survival provide a useful readout of activity and toxicity within this model.

Increased ERK activity is a recognised feature of CRC, most notably in tumours with oncogenic KRAS, NRAS and BRAF (White *et al*., 2026). As expected, all three avatar lines displayed altered ERK activity in the HPZ, with the strongest increase in *byn>RAP-N11*. The data also showed a progressive increase in dpERK signal towards the distal hindgut outside the HPZ, indicating spatial heterogeneity in pathway activity (**Supplementary Figures S2-S4**). Both regorafenib and trametinib reduced ERK activation, although regorafenib produced a more uneven pattern, suggesting incomplete suppression or compensatory pathway reactivation. This aligns with previous studies where MAPK signalling can recover after kinase inhibition through feedback mechanisms (Ponsioen *et al*., 2021). These results show that ERK signalling in these avatars is spatially dynamic and genotype-specific, and this likely contributes to differences in treatment response.

Our redox data provided a second, surprising layer to the avatars’ response to drugs. Oxidative stress contributes to CRC progression, but tumour cells can adapt by increasing antioxidant capacity and reshaping metabolism (Catalano *et al*., 2025). Except for *byn>RAP-N11*, treatment with regorafenib and trametinib generally reduced oxidative stress markers, restored antioxidant measures and improved mitochondrial metabolic balance, consistent with improved fly survival. However, in *byn>RAP-N3* and *byn>RAP-N4*, higher drug concentrations increased mitochondrial metabolic rate, suggesting compensatory metabolic activity under treatment stress. Together with increased *Trx-2* expression in specific settings, this rise points to mitochondrial redox adaptation as a possible mechanism of treatment resistance, in line with previous reports in CRC (Jovanović *et al*., 2022; Qiu *et al*., 2025).

Taken together, these data demonstrate that therapeutic response in the Nigerian patient-derived CRC avatars is strongly shaped by tumour genotype, underlining the limits of a one-size-fits-all approach therapy (Sirugo *et al*., 2019; Vogelstein *et al*., 2013). Further, it opens several questions: which tumours respond to drugs by decreasing mitochondrial metabolic rate, which by elevating it and what are the practical consequences of each tumour type. More broadly, this study strengthens the case for using patient-specific *Drosophila* ‘avatar’ models to test genotype-driven therapeutic effects in colorectal cancer, while addressing an important gap in models representing West African CRC diversity.

## Materials and Methods

### *Drosophila* Models Construction and Validation

The genomic sequence of 10 colorectal cancer patients from existing dataset was accessed through the cBioportal (https://www.cbioportal.org/) (Alatise *et al*., 2021). Cancer driver mutations were selected (see Table 1). To model the effect of loss-of-function mutations in tumour suppressors, shRNA hairpins were created to allow knock-down of the tumour suppressors. A single 21-nucleotide siRNA sequence was designed per gene with DSIR, using a target-adjusted efficiency score of ∼85 (Filhol *et al*., 2012). The target sequences of the siRNAs are given in Supplementary Table S1. Building on our previous work, sense and antisense sequences of designed siRNAs were stitched together into a hairpin cluster, using hairpin-loop-creating and flanking sequences (Bangi *et al*., 2019).

shRNA clusters were synthesised and subcloned into one of the multiple cloning sites (MCSs) of a previously described modified *pWalium* expression vector containing three independent MCSs downstream of UAS response elements (Bangi *et al*., 2019). Synthesis of shRNA clusters and subcloning was performed by GENEWIZ (Azenta Life Sciences). The cDNA transgenes encoding wild-type *Egfr, Pi3K92E* and *Myc* were digested from existing Cagan laboratory vectors and subcloned into the other MCSs as indicated. An example plasmid map is shown in **Supplementary Figure S1**. Prior to injection, the whole plasmid was sequenced by Plasmidsaurus Inc (USA). Plasmid sequences are available on request. PhiC31-mediated transgenic insertion into the VK00018 2^nd^ chromosome landing site was performed by Bestgene. The resulting transgenic lines were crossed to *w^1118^* for two generations to remove the X-chromosomal integrase.

The RAP stock, with an *attP40* insertion of a modified pWalium expression vector carrying UAS-Ras85D^G12V^ and UAS-4X Apc shRNA hairpins + 4X p53 hairpins, was generated previously (Bangi *et al*., 2016; Cong *et al*., 2025). The RA and RP lines (also inserted at *attP40*), were generated in a similar manner to RAP, but lacked p53 or Apc targeting hairpins. Subsequently, the patient-specific transgene was recombined with the RAP transgene to generate the complete model. Presence of the patient-specific (VK00018) and RAP (attP40) insertion in recombinants was confirmed by PCR.

For further validation, transgene expression was targeted in the larvae hindgut using *byn-Gal4*, *Gal80(ts)*, GFP/Tm6b driver line (Cong *et al*., 2025). Virgin female driver flies were crossed with males of patient-specific avatar flies in the ratio of 2:1 after premating for 48 hrs at 25 °C, with *w^1118^* strain as the control. Flies were then allowed to lay eggs for 24 hours on 0.1% dimethyl sulfoxide (DMSO)-containing diets at 18 °C and 27 °C. At 18 °C, embryos were incubated for 24, 48, and 72 hours before being transferred to 27 °C, where development proceeded until eclosion.

### Survival Scoring and Analysis

The embryos from the crossed flies were allowed to develop into pupae and adults, during which they were counted to determine pupation and eclosion, respectively. The eclosion scoring lasted for a period of 7 days post -eclosion, as *byn>transgenes* experienced delay in eclosion. The pupation and eclosion rates were calculated and expressed as percentages of *byn>transgene* (non-tubby) vs. control (*Tb*) animals by the formula outlined below. Calculations: % survival to pupal stage = (EP/CP) × 100, % survival to adult stage = (EA/CP) × 100. [Abbreviations: Control pupae (CP), marked by *Tb*, Experimental pupae (EP), non-tubby and Experimental adults (EA)].

### Model Characterisation and Drug Testing

Crosses were carried out using virgin females of byn-Gal4, Gal80*(ts)*, GFP/Tm6b and male transgene lines on normal diets to pre-mate for 48 hours. Flies were distributed in the ratio 2:1 (female:male) into treatment diets containing either regorafenib or trametinib at final concentrations of 0.01, 0.10 and 1.00 μM, while the control and tumour larvae fed on diet-containing 0.1% DMSO. The flies remained in diets for 24 hours, after which embryos in the diets were allowed to develop at 27 °C. Five (5)-day-old third instar tumour-bearing larvae were collected to perform experiments such as imaging and biochemical assays.

### Antibody Staining and Fluorescent Microscopy

Third instar larvae were placed in pre-washed wipes soaked in 4% sucrose at 27 °C for 2 hours to allow gut clearance and reduce autofluorescence from ingested food (Pranoto & Kwon, 2024). The guts were dissected in ice-cold 1X Phosphate-Buffered Saline (PBS, pH 7.4) and fixed in 4% paraformaldehyde for 30 minutes. Afterwards, the guts were washed 3 times at 5-minute intervals in 1X PBS and then incubated in 1 µg/mL DAPI staining solution (*byn>RAP-N3* and *N11*) and 1 µg/mL of Hoechst in 0.2% PBST (*byn>RAP-N4*) for five minutes. Thereafter, guts were washed 3 times in 1X PBS, after which they were mounted and imaged using Zeiss Axioskop Fluorescent microscope with a PLAN-NEOFLUAR 10X objective and captured using a 5.0 MP 1/2.5’’ APTINA CMOS Digital Camera.

For antibody staining, the gut was dissected and fixed following the method stated above. The fixed gut was washed with PBS plus 0.1% Triton X-100 (0.1% PBST) three times with 15-minute intervals. The gut was then blocked with PBST and Bovine Serum Albumin (PBTB; 1x PBS, 0.1% Triton X-100, 5% BSA (w/v)) for 1 hour at room temperature. The gut was incubated with primary antibody [diphosphor-ERK1/2 (Thr202/Tyr204) rabbit mAb (1:200) (cat no: 9101S)] prepared in 5% PBTB overnight at 4°C. The primary antibody was then washed with 0.1% PBST three times at 15-minute intervals and blocked with PBTB for 1 hour at room temperature. Thereafter, the gut was incubated with the secondary antibody [Alexa Flour 555 goat anti-rabbit IgG (H + L)] at room temperature for 2 hours, followed by Hoechst staining for 5 minutes and washed in 0.1% PBST twice at 5-minute intervals. Finally, guts were mounted with 75% glycerol, and images were visualised using a Zeiss Axioskop Fluorescent microscope with a PLAN-NEOFLUAR 20X objective and captured using a 5.0 MP 1/2.5’’ APTINA CMOS Digital Camera. The fluorescent intensity was obtained using Fiji software.

### Image Analysis for Fluorescent Intensity

After imaging the guts using Image view software vx64,3.7.6701, fluorescent signal intensity was quantified using Fiji software (Schindelin *et al*., 2012). Green channel images of the HPZ were separated and background subtracted at 200 before quantification. The region of interest for GFP measurement was manually marked using the *polygon* selection tool encompassing the HPZ to the luminal area of the hindgut and measured for the GFP intensity. Similarly, the ROI for dpERK signal intensity was also selected with the *oval* tool, set around the posterior midgut-hindgut junction. The dpERK spatial intensity profile (**Supplementary Figures S2-S4**) was measured using the 140 µm line manually drawn from the HPZ to the distal part of the hindgut. Data obtained from Fiji was analysed using GraphPad Prism while dpERK intensity profile was visualised using Microsoft Power BI Desktop, Version 2.152.1057.0 64-bit (March 2026).

### Larvae Imaging

Wandering larvae were immobilised in PBS for 3-4 hours at 4°C, followed by imaging and capturing using a 5.0MP 1/2.5’’ APTINA CMOS Digital Camera attached to a dissecting light microscope at a magnification of 0.7X.

### Larvae and Relative Gut Size Measurement and Analysis

HPZ and larval areas were measured using Fiji (ImageJ v1.54p). Gut images were acquired at 10X magnification using a Zeiss Axioskop fluorescent microscope with a PLAN-NEOFLUAR 10X objective, and larvae were imaged at 0.7X using StereoBlue Euromex stereo microscope. A 5.0 MP 1/2.5ʺ Aptina CMOS digital camera was utilised for image collection. Images were calibrated to millimetres based on pixel dimensions of the camera (2592 × 1944 with a pixel pitch of ≈ 2.2 µm/pixel).

To measure larval length and width, FiJi was scaled to 3.143 µm/pixel after adjustments for magnification (µm/pixel = 2.2 / 0.7 = 3.143 µm/pixel). Relative gut size was quantified using the dimensions of the hindgut proliferation zone following a similar calibration approach used in larval measurements with adjustment for magnification (µm/pixel = 2.2 / 10 = 0.22 µm/pixel). The length of the HPZ to luminal area was measured, while its width was measured at three distinct regions along its span. The relative gut size was then determined by normalising gut area to the overall larva size (area) using the formula: Relative gut size = Area of the gut / Area of the larvae size. The normalisation accounts for the variation in larva body size and allows for accurate comparison of the gut across samples.

### Preparation of *D. melanogaster* Larvae for Biochemical Assays

To perform biochemical assays, the larvae were weighed using ES 225SM-DR SWISS MADE weighing balance and homogenised in cold 0.1 M phosphate buffer, pH 7.4 (10 μL per mg), on ice. The homogenate was centrifuged using Eppendorf centrifuge 5804 R at 2300 rcf for 15 minutes at 4 °C. The obtained supernatant was gently pipetted into sterile microfuge tubes and utilised for the examination of biochemical markers of oxidative stress and antioxidants.

### Protein Determination

Protein concentration was determined using Lowry method (Kielkopf *et al*., 2020). In summary, the reaction mixture contained 20 µL of diluted sample (1:10 dilution), 60 µL of distilled water and 200 µL of solution C (0.1 M NaOH, 2% Na_2_CO_3_, 2% Na-K tartrate and 1% CuSO_4_). The reaction mixture was incubated at room temperature for 15 minutes, after which 20 μL of 1:5 dilutions of Folin-Ciocalteu reagent was added and incubated for another 15 minutes. Absorbance was measured using a SpectraMax Plus 384 microplate reader (Molecular Devices, San Jose, CA, USA) at 650 nm against a blank. Protein determination assay was carried out to normalise total thiol and non-protein thiol contents.

### Estimation of Total Thiols Level

Total thiol concentration was determined using Ellman reagent (Dhawan *et al*., 2021). The reaction mixture consists of 20 µL of the prepared diluted sample, 10 µL of 5,5′-dithiobis (2-nitrobenzoic acid) (DTNB) and 270 µL of 0.1 M phosphate buffer. The reaction mixture was incubated for 30 minutes at room temperature, and absorbance was taken at 412 nm using SpectraMax Plus 384 microplate reader (Molecular Devices, San Jose, CA, USA). the total thiol level was calculated using GSH standard curve and expressed as µmol/mg protein.

### Determination of Non-Protein Thiol Level

Non-Protein Thiol (NPSH)_level was estimated using Ellman’s reagent. The protein content of the sample was precipitated using 10% Trichloroacetic Acid (TCA) in the ratio 1:1. Samples were kept at 4 °C for 1 hr, then centrifuged at 5000 rpm for 10 min at 4 °C. The reaction mixture consisted of 200 µL of 0.1 M phosphate buffer, 50 µL of sample, and 50 µL of DTNB. The absorbance was read at 412 nm using SpectraMax Plus 384 microplate reader (Molecular Devices, San Jose, CA, USA). The concentration of the NPSH was estimated using a standard curve and presented as mmol/mg of protein (Ginet *et al*., 2025).

### ROS/RONS Oxidation Assessment

The oxidation of 2’,7’-dichlorofluorescein (DCFH) by the sample supernatant serves as a broad index of oxidative stress, as previously described (Pérez-Severiano *et al*., 2004). The reaction mixture consisted of 450 µL of 0.1 µM potassium phosphate buffer (pH 7.4), 120 µL of distilled water, 15 µL of 200 µM DCFH-DA (final concentration: 5 µM), and 15 µL of diluted sample. The fluorescence emission of DCF produced by DCFH oxidation was recorded for 10 minutes (every 30 seconds) at excitation and emission wavelengths of 488 nm and 525 nm, respectively, using a fluorescence spectrophotometer (Infitek SP-LF96S). The degree of DCF formation was represented as a percentage of the experimental control group.

### Estimation of Nitrite/Nitrate (Nitric oxide) Level

The nitrite/nitrate (nitric oxide) concentration was determined according to the Green method (Tsikas, 2007). The reaction mixture contained equal volume of freshly prepared Griess reagent (0.1% N-(1-naphthyl) ethylenediamine dihydrochloride; 1% sulfanilamide in phosphoric acid) and the prepared sample (1:1), which was incubated, and the absorbance was measured at 550 nm using SpectraMax Plus 384 microplate reader (Molecular Devices, San Jose, CA, USA). Nitric oxide level was quantified using the sodium nitrate standard curve measured at 550 nm.

### Estimation of the Mitochondrial Metabolic Rate

Mitochondrial metabolic rate of the whole treated larvae was determined using MTT assay (Abe & Matsuki, 2000). Reaction mixture containing 20 μL of diluted sample, 10 μL of 3-(4,5-dimethylthiazol-2-yl)-2,5-diphenyltetrazolium bromide (MTT) at a final concentration of 5 mg/mL and 170 μL of PBS was incubated at 37 °C for 3 hours. The formazan product was then read using SpectraMax Plus 384 microplate reader (Molecular Devices, San Jose, CA, USA) at 570 nm and 650 nm after dissolution in 100 μL of DMSO. Mitochondrial metabolic rate was represented as percentage of control group.

### RNA Extraction and Analysis

Total RNA was isolated from 10-15 dissected third instar larvae guts of CRC avatars using the phenol-chloroform method described previously (Green & Sambrook, 2020). The purity of the isolated RNA was checked at A260/A280 and A260/A230 using the Spectradrop micro-volume microplate in SpectraMax Plus 384 microplate reader (Molecular Devices, San Jose, CA, USA).

### Primer Design and Efficiency

The genes used in this study were obtained from (https://flybase.org/). Primers were designed using Primer3 (https://primer3.ut.ee/) and the primer BLAST tool of the NCBI (https://www.ncbi.nlm.nih.gov/tools/primer-blast/). Primer Sequence for *Trx-2* Forward Primer (GTCTCCACATCTCCCATCCA) and *Trx-2* Reverse Primer (TTGACGCCGTTCTTGAGGAA). To ensure specificity and precise amplification of the target genes, the designed primer was subjected to validation as reported previously (Chen *et al*., 2023).

### cDNA Synthesis and qRT-PCR Analysis

An aliquot of the isolated total RNA (200 ng) was used for the cDNA synthesis following the manufacturer’s described protocol for the LunaScript® RT SuperMix Kit (E3010) and carried out in a BioRad T100 conventional PCR machine. The Luna Universal qPCR master mix kit was used for the qRT-PCR. For each of the replicates, a 10 µL total reaction volume was used, consisting of 1.5 µL of cDNA from each sample and 8.5 µL of PCR reaction mix (5 µL TaqMan master mix, 3 µL nuclease-free water and 0.5 µL gene expression assay primer-probe mix). Standard fast thermal cycling parameters of 40 cycles of 95 °C for 15 seconds and 55 °C for 45 seconds were applied in accordance with the manufacturer’s recommendations using q-PCR thermocycler QuantStudio-5. The relative quantities were corrected for efficiency of amplification, and fold change in gene expression between groups was calculated using the comparative quantification method (Pfaffl, 2001).

## Statistical Analysis

Data are presented as median or mean ± SEM, or range as indicated. Comparisons among multiple groups or factors were carried out using one-way or two-way ANOVA, followed by Tukey’s post hoc test for multiple comparisons. Two-tailed unpaired Student’s t-test was used for comparisons between two independent groups. Pearson’s correlation analysis was conducted to assess linear relationships between continuous variables. Parametric test assumptions were verified before analysis. A *p* < 0.05 was considered statistically significant. Analyses were carried out using GraphPad Prism version 9.0.0.

---

**Supplemental Table 1.**
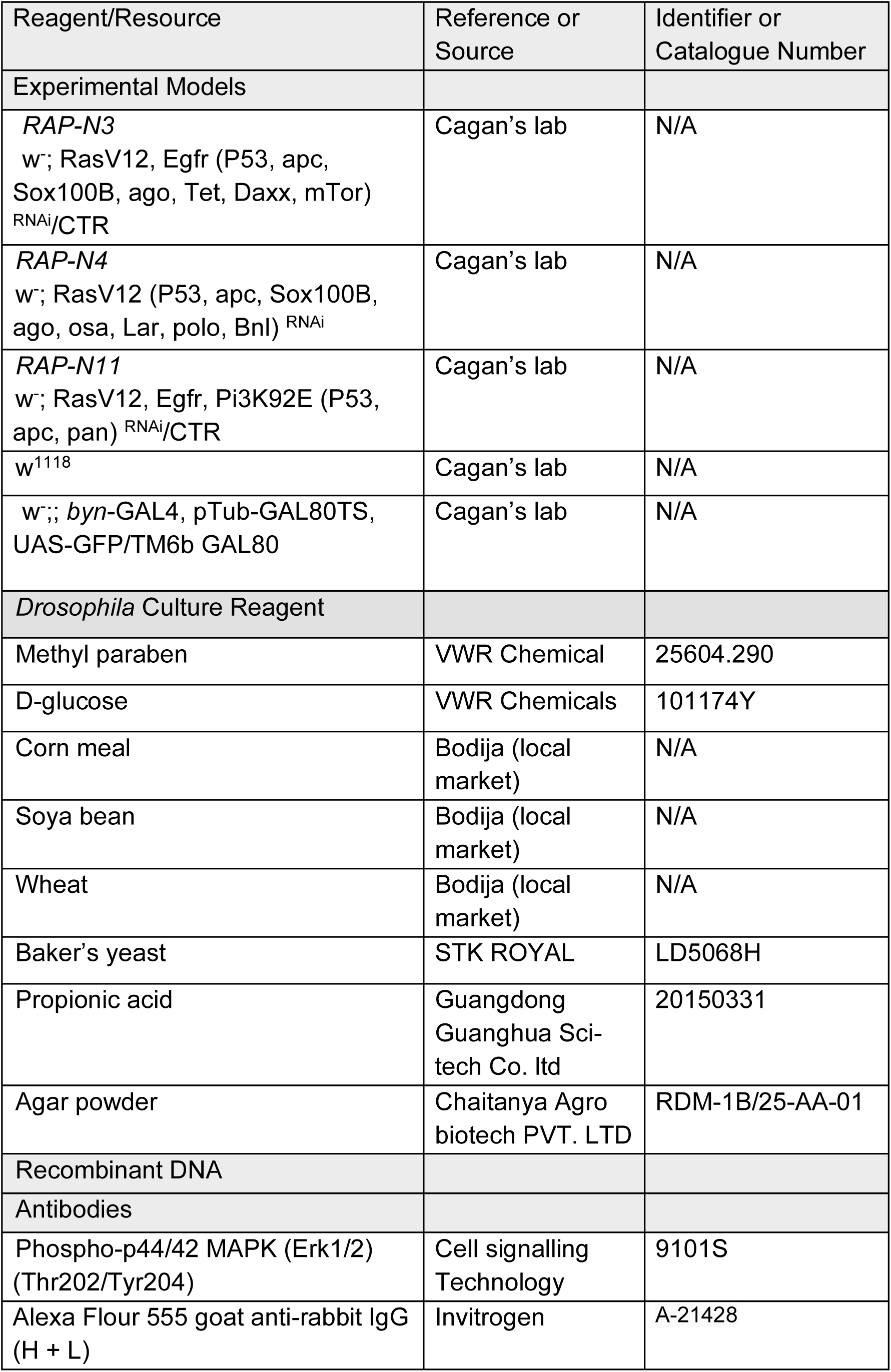
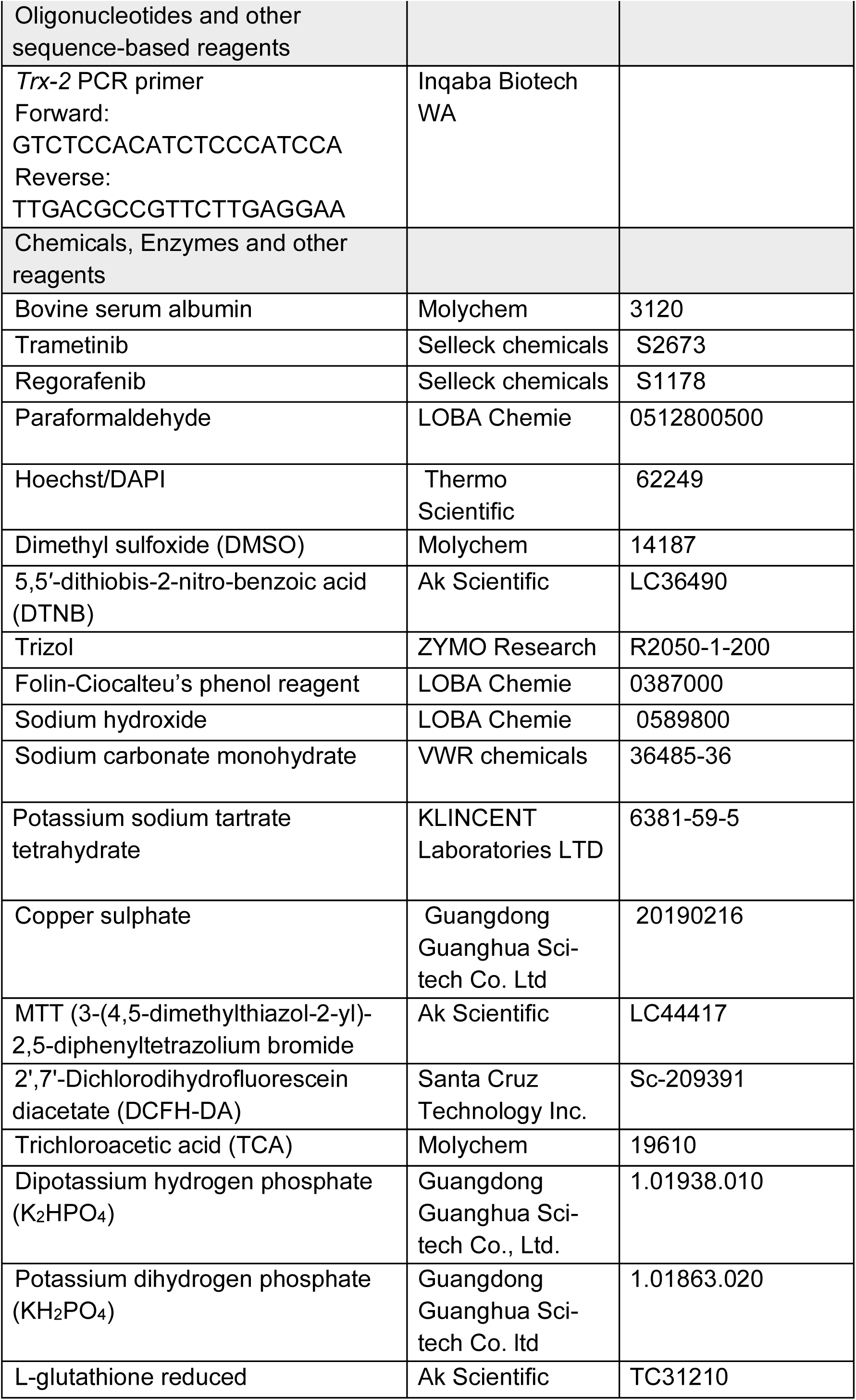
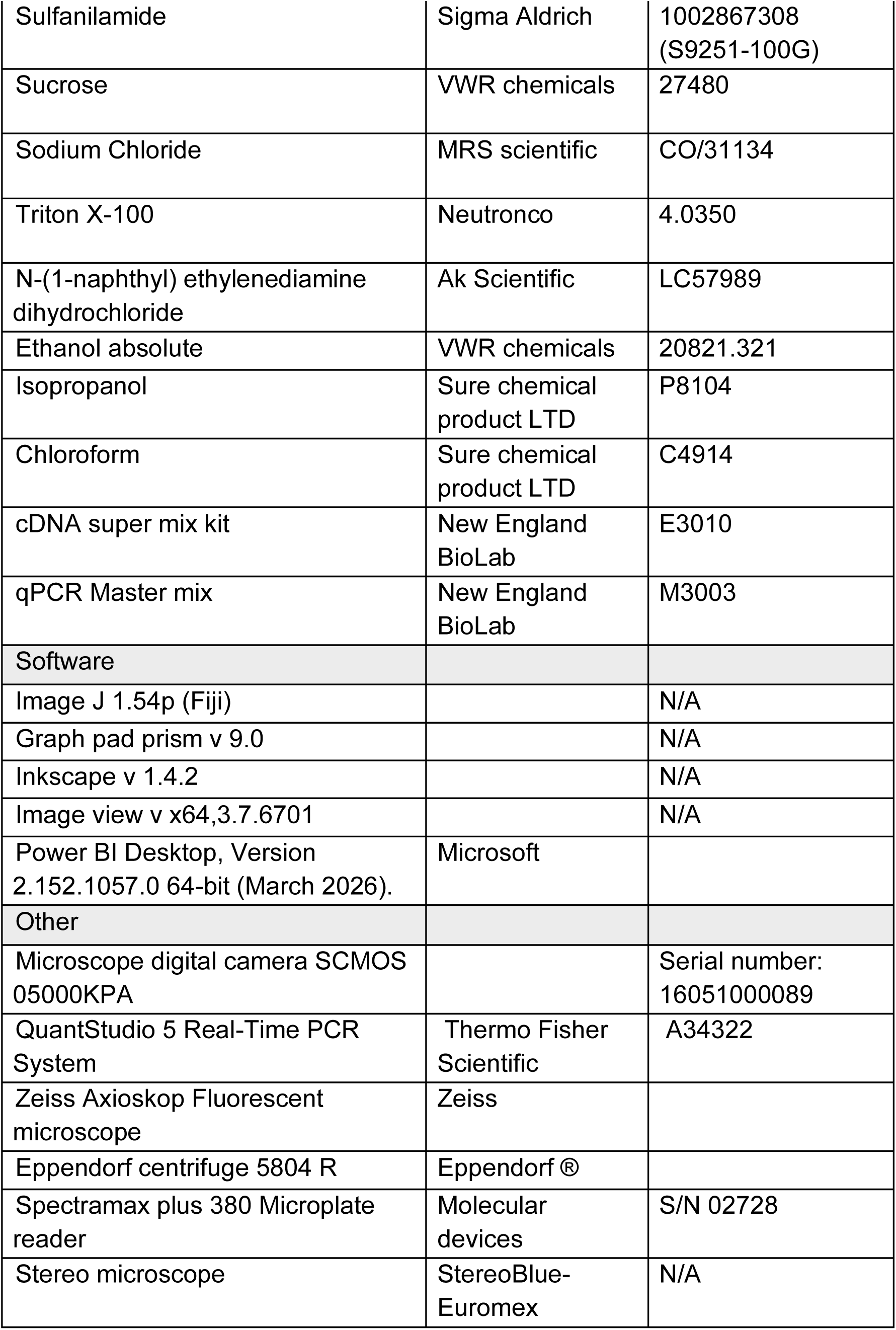
Reagents and tools.

## Acknowledgements

We thank the members of our laboratories for their helpful discussions. The three CRC *Drosophila melanogaster* models generated in this study were derived from genomic datasets deposited in cBioPortal by Alatise and colleagues (Alatise *et al*., 2021), whose contributions we gratefully acknowledge. R.C. gratefully acknowledges support from the NIH (R01CA258736), a Royal Society Wolfson Fellowship, Chief Scientific Office (EPD/22/13) (TCS/22/02), CRUK (CTRQQR-2021\100006) Pershing Square Sohn, and Baillie-Gifford to R.C. H.A.B. gratefully acknowledges support by Tenovus Scotland (S23-01) and a Scotland CSO Fellowship (EPD/23/01).

## Authors’ Contributions

**Festus Adeyimika Oyeniyi:** Investigation; Formal analysis; Visualisation; Writing-Original Draft; **Favour Ayomide Oladokun:** Investigation; Formal analysis; Visualisation; Writing-Original Draft; **Abigael Abosede Ajayi:** Investigation; Formal analysis; Visualisation; Writing-Original Draft; **Abdulwasiu Ibrahim:** Formal analysis; Writing-Original Draft; **Ruth Seyi Aladeloye:** Investigation; Writing-Original Draft; **Opeyemi Abigail Akinfe**: Investigation; Writing-Original Draft; **Funmilayo Rosemary Oludaiye:** Investigation; Writing-Original Draft; **Thomas G. Moens:** Model building, Methodology; review and editing; **Hammed A. Badmos:** Supervision; Model building; Methodology; review and editing. **Amos Olalekan Abolaji:** Supervision; Methodology; Model building, Funding acquisition; review and editing. **Ross Cagan:** Conceptualisation; Supervision; Model building, Methodology; Funding acquisition; Review and editing.

## Data Availability

All the data supporting the findings of this study are provided in the main text and Supplementary Materials. Any additional information or datasets that could further aid in understanding this work are available from the corresponding authors upon reasonable request.

## Ethics Declarations

The authors declare no competing interests.

## Supplementary Information

**Table S1:**
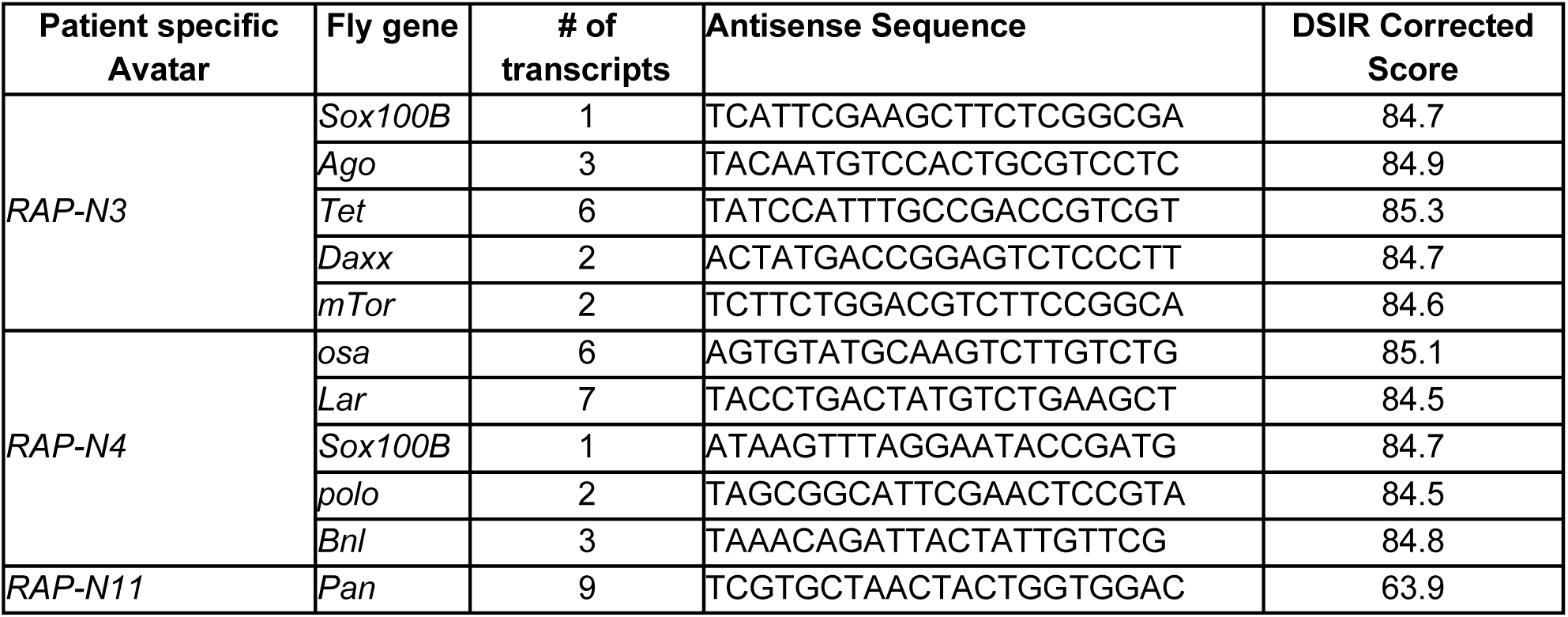
Sequences of single 21-nucleotide siRNA for knockdown of tumour suppressors.

**Table S2:**
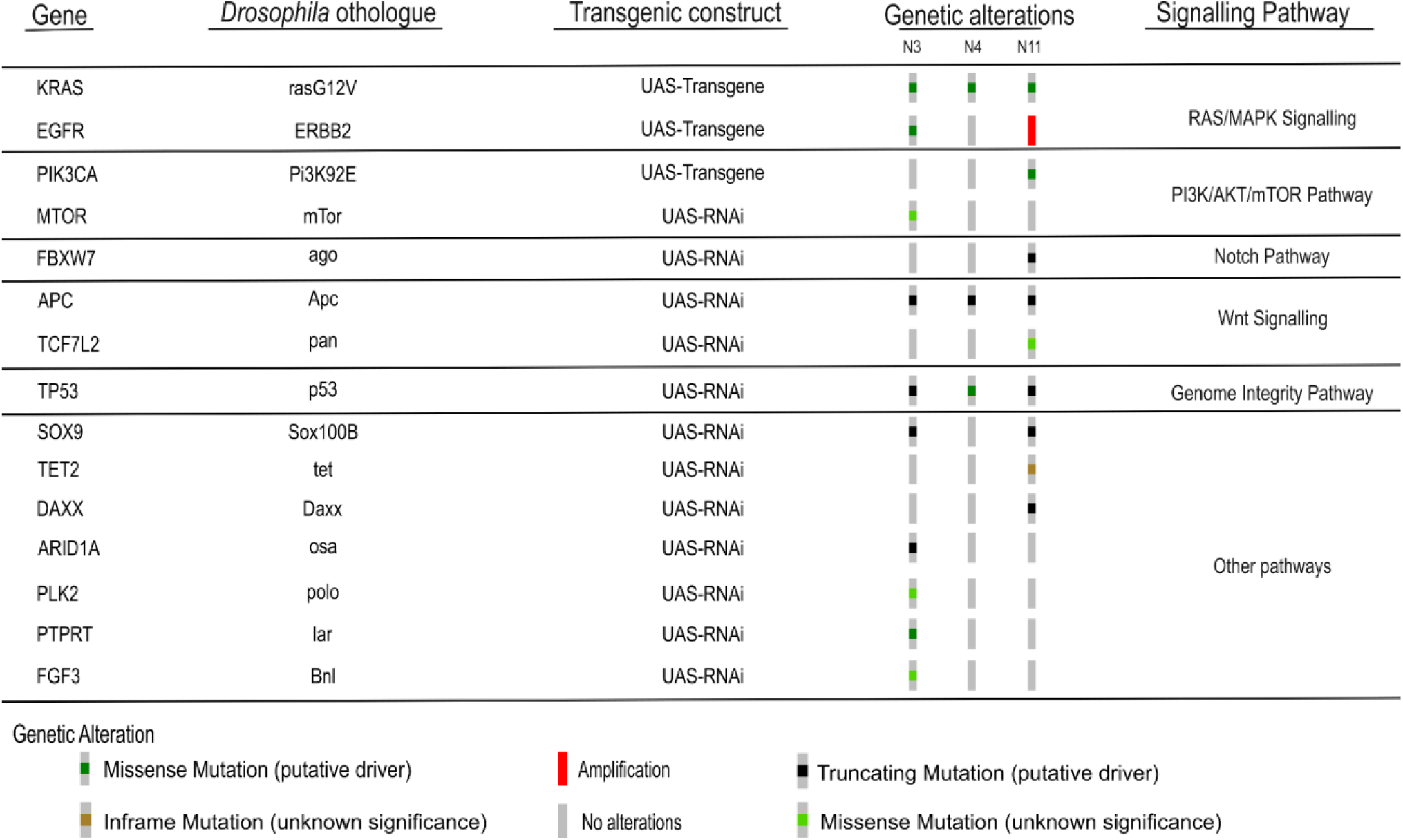
Selected Nigerian patient-specific colorectal cancer driver genes, their *Drosophila* orthologues, and associated signalling pathways.

**Table S3:** Summary of the selected three patient-derived colorectal cancer Fly avatars and their genetic alterations.

| Avatar ID | Patient ID | Base | No. of mutations | shRNAs | Oncogenes |
| --- | --- | --- | --- | --- | --- |
| N3 | pkc_crc_083 | RAP | 9 | *Sox100B* (SOX9), *ago* (FBXW7), *Tet* (TET2), *Daxx* (DAXX), *mTor* (MTOR) | *Egfr* (ERBB2) |
| N4 | pkc_crc_137 | RAP | 8 | *osa* (ARID1A), *Sox 100B* (SOX9), *Lar* (PTPRT), *polo* (PLK2), *Bnl* (FGF3) |  |
| N11 | pkc_crc_079 | RAP | 6 | *pan* (TCF7L2) | *Pi3K92E* (PIK3CA), *Egfr* (ERBB2) |

**Figure S1:**
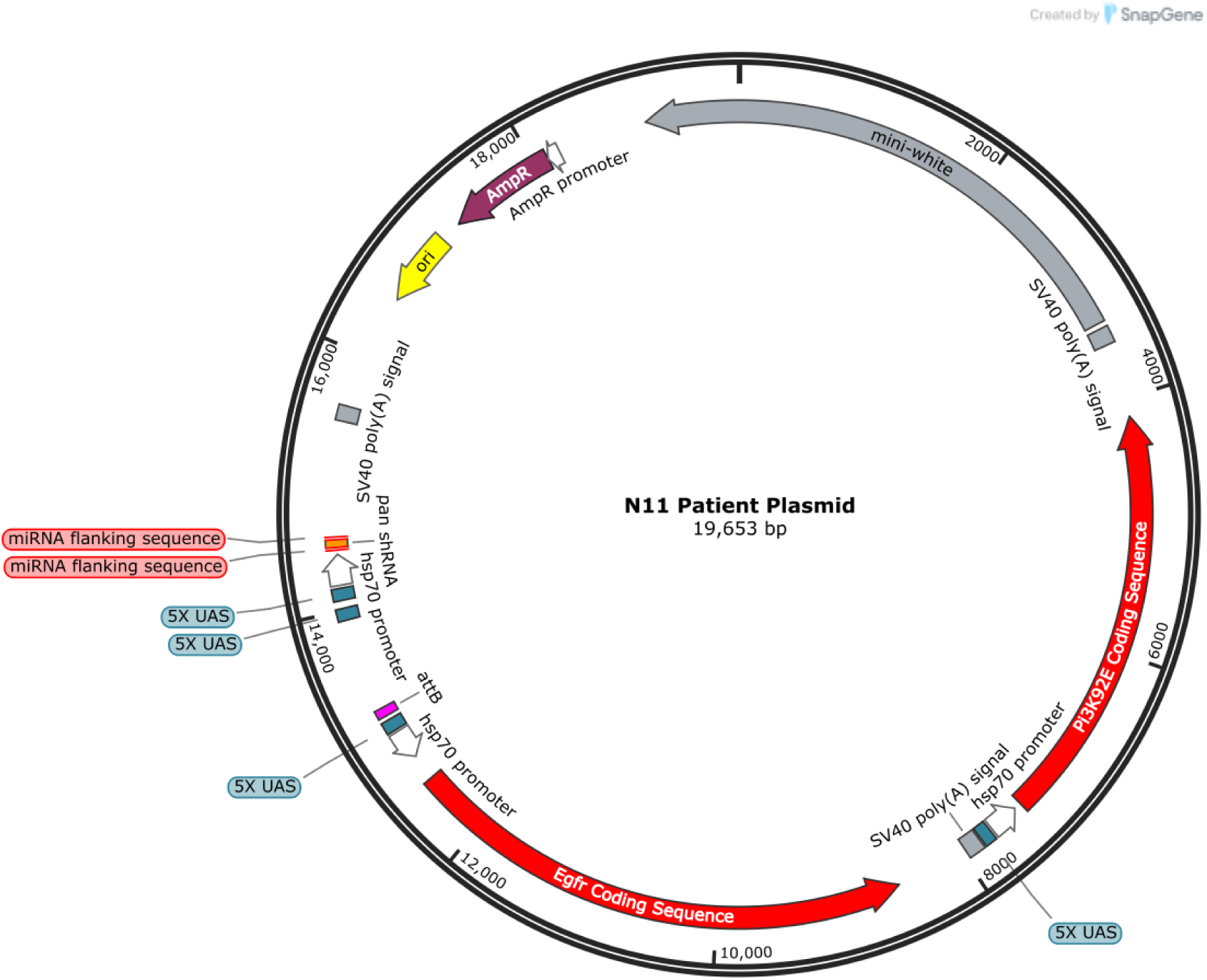
Cloning vector for transgenesis

**Figure S2:**
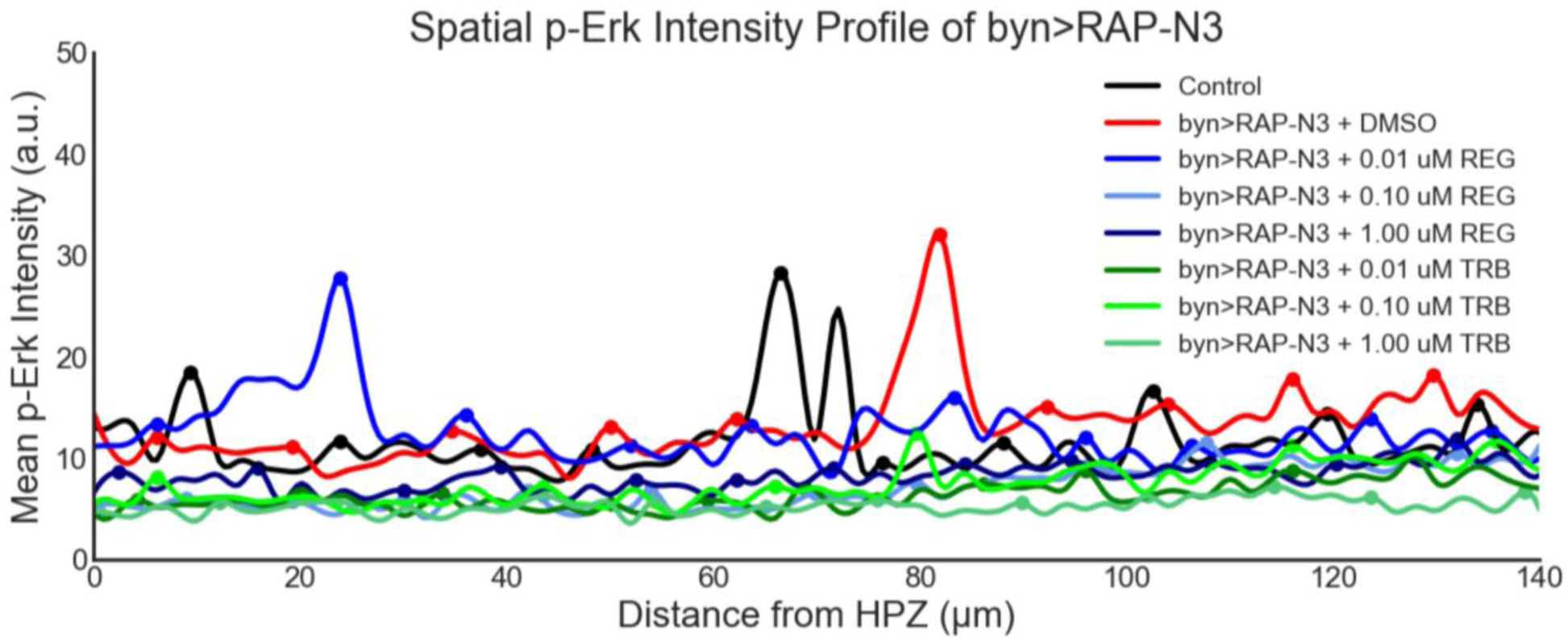
Spatial Intensity profile of phospho-ERK in regorafenib and trametinib fed *byn > RAP-N3*

**Figure S3:**
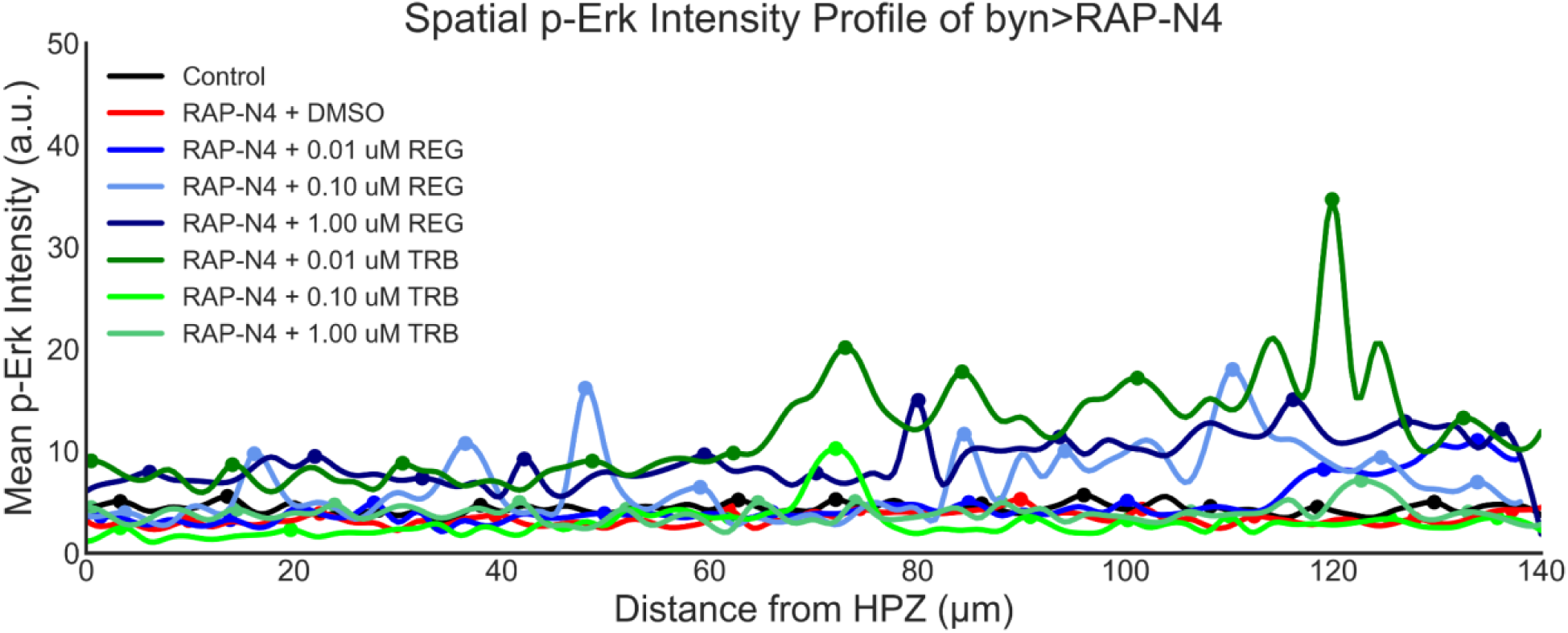
Spatial Intensity profile of phospho-ERK in regorafenib and trametinib fed *byn > RAP-N4*

**Figure S4:**
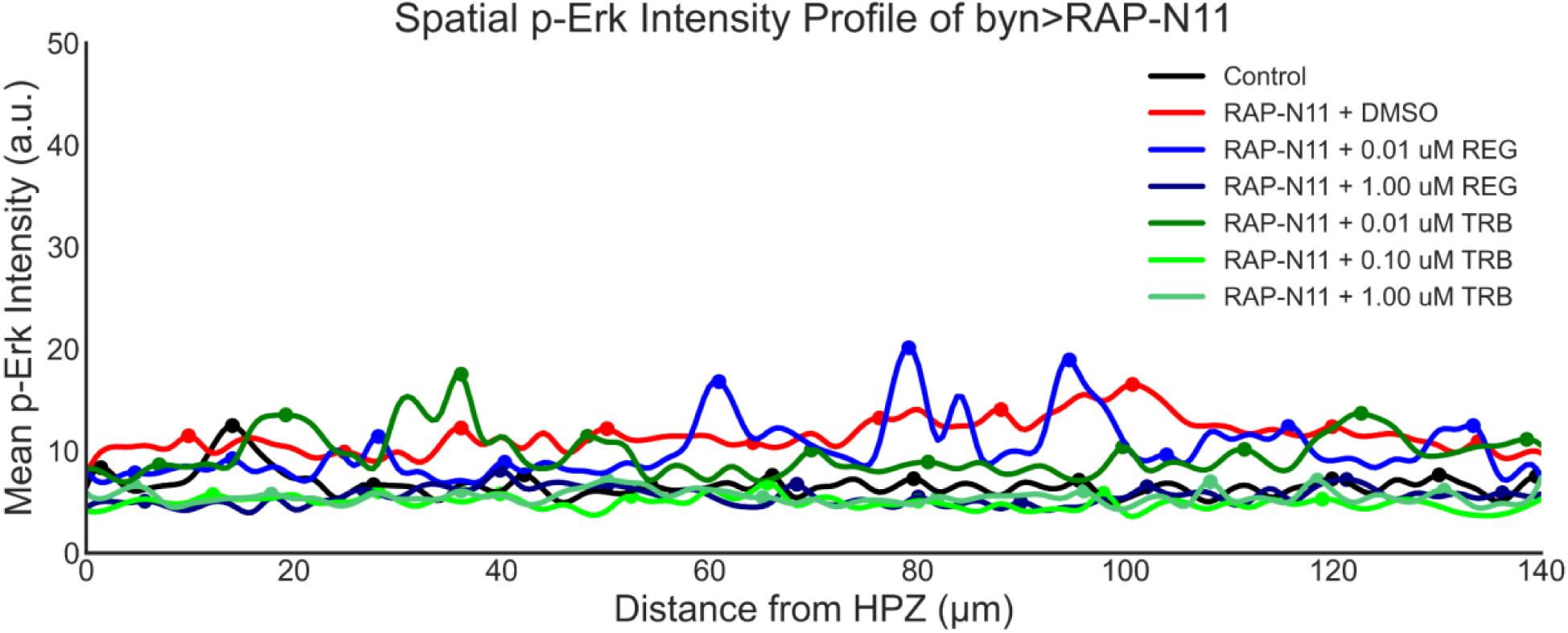
Spatial Intensity profiles of phospho-ERK in regorafenib and trametinib fed *byn > RAP-N11*

### Note S1

Fly crosses

Crosses were done using virgin females of *byn*-*Gal4*, *Gal80(ts)*, *GFP*/*TM6b*, and male Transgene lines in the ratio 2:1. Flies were pre-mated for 48 hours at 27 ℃ before the various groupings. The experiment was carried out at 27 ℃, and the adult flies were moved out of the drug-containing diet after 24 hours of egg laying. Third instar larvae were collected for further studies.

### Note S2

Temperature calibration

To determine the most appropriate condition for the fly model under study, temperature calibration of the Nigerian CRC avatar lines was carried out in different temperature conditions. Flies were pre-mated for 48 hours at 27℃ before the various groupings.

Four different conditions for calibration.

1. The flies were left to mate for 24 hours at 27 °C, after which the embryo-containing diet was left at 27 °C until the third instar larva emerged.
2. The flies were allowed to lay for 24 hours at 18 °C, then the embryo-containing diet was moved to 27 °C until the third instar larva emerged.
3. The flies were allowed to lay for 24 hours at 18 °C. The embryo-containing diet was left at 18 °C for an additional 24 hours and then transferred to 27 °C until the third-instar larvae emerged.
4. The flies were allowed to lay for 24 hours at 18 °C. The embryo-containing diet was left at 18 °C for an additional 48 hours and then moved to 27 °C until the third instar larva emerge.

### Correlation Analysis for Relative gut size vs other biomarkers

**Table S4:** Correlation analysis of gut size and assayed biomarkers of regorafenib-treated *byn>RAP-N3* fly avatar.

|  | Relative Gut Size vs. ECLOSION | Relative Gut Size vs. TSH | Relative Gut Size vs. NPSH | Relative Gut Size vs. NO | Relative Gut Size vs. RONS | Relative Gut Size vs. MTT | Relative Gut Size vs. GFP Intensity |
| --- | --- | --- | --- | --- | --- | --- | --- |
| Pearson r |  |  |  |  |  |  |  |
| r | -0.9910 | 0.1832 | -0.2116 | 0.2791 | 0.1356 | 0.02199 | -0.8312 |
| 95% confidence interval | -0.9998 to -0.6279 | -0.9441 to 0.9730 | -0.9745 to 0.9408 | -0.9320 to 0.9779 | -0.9492 to 0.9702 | -0.9594 to 0.9627 | -0.9963 to 0.6458 |
| R squared | 0.9820 | 0.03357 | 0.04475 | 0.07788 | 0.01839 | 0.0004835 | 0.6908 |
| P value |  |  |  |  |  |  |  |
| P (two-tailed) | 0.009 | 0.82 | 0.79 | 0.72 | 0.86 | 0.98 | 0.17 |
| P-value summary | ** | ns | ns | ns | ns | ns | ns |
| Significant? (alpha = 0.05) | Yes | No | No | No | No | No | No |

**Table S5:** Correlation analysis of gut size and assayed biomarkers of trametinib-treated *byn>RAP-N3* fly avatar.

|  | Relative Gut<br>Size<br>vs.<br>ECLOSION | Relative Gut<br>Size<br>vs.<br>TSH | Relative Gut<br>Size<br>vs.<br>NPSH | Relative Gut<br>Size<br>vs.<br>NO | Relative Gut<br>Size<br>vs.<br>RONS | Relative Gut<br>Size<br>vs.<br>MTT | Relative Gut<br>Size<br>vs.<br>GFP Intensity |
| --- | --- | --- | --- | --- | --- | --- | --- |
| Pearson r |  |  |  |  |  |  |  |
| r | -0.9562 | 0.5531 | -0.1740 | 0.2269 | 0.07643 | -0.5157 | -0.9169 |
| 95% confidence<br>interval | -0.9991 to<br>0.06051 | -0.8710 to<br>0.9886 | -0.9725 to<br>0.9451 | -0.9389 to<br>0.9753 | -0.9548 to<br>0.9665 | -0.9874 to<br>0.8831 | -0.9983 to<br>0.3722 |
| R squared | 0.9143 | 0.3059 | 0.03028 | 0.05148 | 0.005842 | 0.2659 | 0.8406 |
| P value |  |  |  |  |  |  |  |
| P (two-tailed) | 0.04 | 0.45 | 0.83 | 0.77 | 0.92 | 0.48 | 0.08 |
| P value summary | * | ns | ns | ns | ns | ns | ns |
| Significant?<br>(alpha = 0.05) | Yes | No | No | No | No | No | No |

**Table S6:** Correlation analysis of gut size and assayed biomarkers of regorafenib-treated *byn>RAP-N4* fly avatar.

|  | Relative Gut Size vs. ECLOSION | Relative Gut Size vs. TSH | Relative Gut Size vs. NPSH | Relative Gut Size vs. NO | Relative Gut Size vs. RONS | Relative Gut Size vs. MTT | Relative Gut Size vs. GFP Intensity |
| --- | --- | --- | --- | --- | --- | --- | --- |
| Pearson r |  |  |  |  |  |  |  |
| r | -0.7130 | 0.8050 | 0.3293 | -0.9580 | -0.7911 | -0.8756 | -0.8500 |
| 95% confidence interval | -0.9934 to 0.7882 | -0.6896 to 0.9957 | -0.9243 to 0.9802 | -0.9991 to 0.03926 | -0.9954 to 0.7093 | -0.9974 to 0.5395 | -0.9968 to 0.6068 |
| R squared | 0.5083 | 0.6481 | 0.1084 | 0.9177 | 0.6258 | 0.7666 | 0.7225 |
| P value |  |  |  |  |  |  |  |
| P (two-tailed) | 0.29 | 0.19 | 0.67 | 0.04 | 0.21 | 0.12 | 0.15 |
| Significant? (alpha = 0.05) | No | No | No | Yes | No | No | No |

**Table S7:**
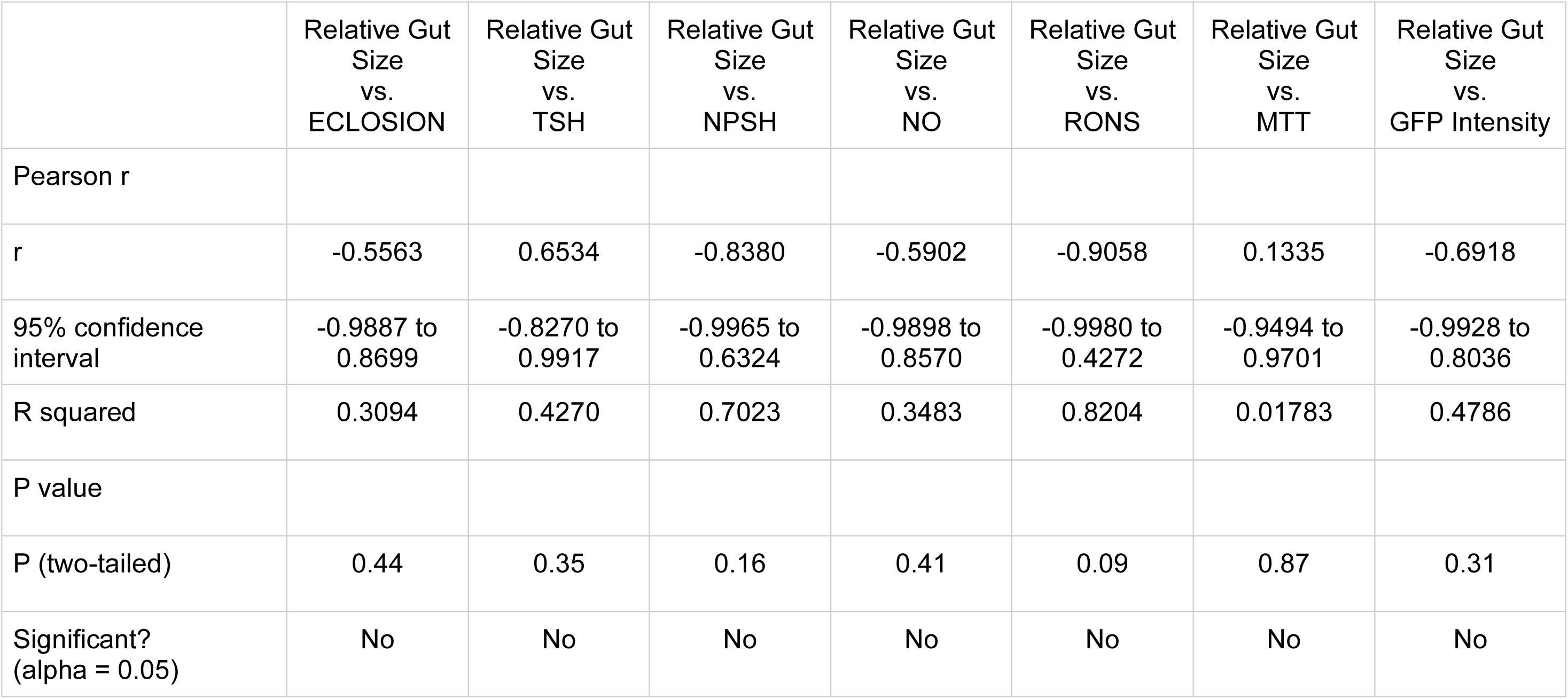
Correlation analysis of gut size and assayed biomarkers of trametinib-treated *byn>RAP-N4* fly avatar.

**Table S8:** Correlation analysis of relative gut size and assayed biomarkers of regorafenib-treated *byn>RAP-N11* fly avatar.

|  | Relative Gut Size vs. Eclosion | Relative Gut Size vs. TSH | Relative Gut Size vs. NPSH | Relative Gut Size vs. NO | Relative Gut Size vs. RONS | Relative Gut Size vs. MTT | Relative Gut Size vs. GFP Intensity |
| --- | --- | --- | --- | --- | --- | --- | --- |
| Pearson r |  |  |  |  |  |  |  |
| r | -0.9326 | 0.8716 | 0.8378 | -0.8423 | 0.1194 | 0.09112 | -0.5165 |
| 95% confidence interval | -0.9986 to 0.2750 | -0.5514 to 0.9973 | -0.6329 to 0.9965 | -0.9966 to 0.6237 | -0.9508 to 0.9693 | -0.9535 to 0.9675 | -0.9874 to 0.8828 |
| R squared | 0.8697 | 0.7597 | 0.7019 | 0.7095 | 0.01427 | 0.008303 | 0.2668 |
| P value |  |  |  |  |  |  |  |
| P (two-tailed) | .067 | .128 | .162 | .158 | .881 | .909 | .483 |
| Significant? (alpha = 0.05) | No | No | No | No | No | No | No |

**Table S9:**
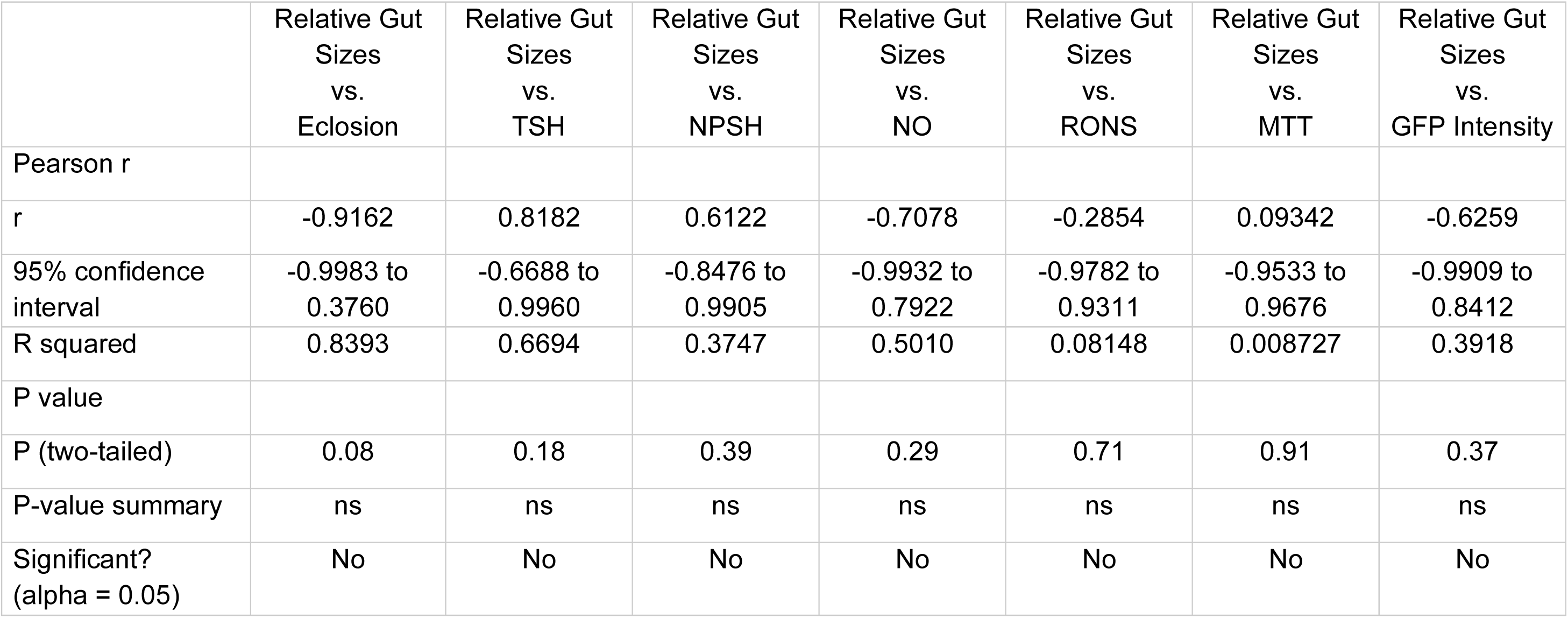
Correlation analysis of gut size and assayed biomarkers of trametinib-treated *byn>RAP-N11* fly avatar.

|  | Relative Gut<br>Sizes<br>vs.<br>Eclosion | Relative Gut<br>Sizes<br>vs.<br>TSH | Relative Gut<br>Sizes<br>vs.<br>NPSH | Relative Gut<br>Sizes<br>vs.<br>NO | Relative Gut<br>Sizes<br>vs.<br>RONS | Relative Gut<br>Sizes<br>vs.<br>MTT | Relative Gut<br>Sizes<br>vs.<br>GFP Intensity |
| --- | --- | --- | --- | --- | --- | --- | --- |
| Pearson r |  |  |  |  |  |  |  |
| r | -0.9162 | 0.8182 | 0.6122 | -0.7078 | -0.2854 | 0.09342 | -0.6259 |
| 95% confidence<br>interval | -0.9983 to<br>0.3760 | -0.6688 to<br>0.9960 | -0.8476 to<br>0.9905 | -0.9932 to<br>0.7922 | -0.9782 to<br>0.9311 | -0.9533 to<br>0.9676 | -0.9909 to<br>0.8412 |
| R squared | 0.8393 | 0.6694 | 0.3747 | 0.5010 | 0.08148 | 0.008727 | 0.3918 |
| P value |  |  |  |  |  |  |  |
| P (two-tailed) | 0.08 | 0.18 | 0.39 | 0.29 | 0.71 | 0.91 | 0.37 |
| P-value summary | ns | ns | ns | ns | ns | ns | ns |
| Significant?<br>(alpha = 0.05) | No | No | No | No | No | No | No |

